# High Frequencies of PD-1^+^TIM3^+^TIGIT^+^CTLA4^+^ Functionally Exhausted SARS-CoV-2-Specific CD4^+^ and CD8^+^ T Cells Associated with Severe Disease in Critically ill COVID-19 Patients

**DOI:** 10.1101/2022.01.30.478343

**Authors:** Pierre-Gregoire Coulon, Swayam Prakash, Nisha R. Dhanushkodi, Ruchi Srivastava, Latifa Zayou, Delia F. Tifrea, Robert A. Edwards, J. Figueroa Cesar, Sebastian D. Schubl, Lanny Hsieh, Anthony B. Nesburn, Baruch D. Kuppermann, Elmostafa Bahraoui, Hawa Vahed, Daniel Gil, Trevor M. Jones, Jeffrey B. Ulmer, Lbachir BenMohamed

## Abstract

SARS-CoV-2-specific memory T cells that cross-react with common cold coronaviruses (CCCs) are present in both healthy donors and COVID-19 patients. However, whether these cross-reactive T cells play a role in COVID-19 pathogenesis versus protection remain to be fully elucidated. In this study, we characterized cross-reactive SARS-CoV-2-specific CD4^+^ and CD8^+^ T cells, targeting genome-wide conserved epitopes in a cohort of 147 non-vaccinated COVID-19 patients, divided into six groups based on the degrees of disease severity. We compared the frequency, phenotype, and function of these SARS-CoV-2-specific CD4^+^ and CD8^+^ T cells between severely ill and asymptomatic COVID-19 patients and correlated this with α-CCCs and β-CCCs co-infection status. Compared with asymptomatic COVID-19 patients, the severely ill COVID-19 patients and patients with fatal outcomes: (*i*) Presented a broad leukocytosis and a broad CD4^+^ and CD8^+^ T cell lymphopenia; (*ii*) Developed low frequencies of functional IFN-*γ*-producing CD134^+^CD138^+^CD4^+^ and CD134^+^CD138^+^CD8^+^ T cells directed toward conserved epitopes from structural, non-structural and regulatory SARS-CoV-2 proteins; (*iii*) Displayed high frequencies of SARS-CoV-2-specific functionally exhausted PD-1^+^TIM3^+^TIGIT^+^CTLA4^+^CD4^+^ and PD-1^+^TIM3^+^TIGIT^+^CTLA4^+^CD8^+^ T cells; and (*iv*) Displayed similar frequencies of co-infections with β-CCCs strains but significantly fewer co-infections with α-CCCs strains. Interestingly, the cross-reactive SARS-CoV-2 epitopes that recalled the strongest CD4^+^ and CD8^+^ T cell responses in unexposed healthy donors (HD) were the most strongly associated with better disease outcome seen in asymptomatic COVID-19 patients. Our results demonstrate that, the critically ill COVID-19 patients displayed fewer co-infection with α-CCCs strain, presented broad T cell lymphopenia and higher frequencies of cross-reactive exhausted SARS-CoV-2-specific CD4^+^ and CD8^+^ T cells. In contrast, the asymptomatic COVID-19 patients, appeared to present more co-infections with α-CCCs strains, associated with higher frequencies of functional cross-reactive SARS-CoV-2-specific CD4^+^ and CD8^+^ T cells. These findings support the development of broadly protective, T-cell-based, multi-antigen universal pan-Coronavirus vaccines.

**KEY POINTS:**

- A broad lymphopenia and lower frequencies of SARS-CoV-2-specific CD4^+^ and CD8^+^ T-cells were associated with severe disease onset in COVID-19 patients.
- High frequencies of phenotypically and functionally exhausted SARS-CoV-2-specific CD4^+^ and CD8^+^ T cells, co-expressing multiple exhaustion markers, and targeting multiple structural, non-structural, and regulatory SARS-CoV-2 protein antigens, were detected in severely ill COVID-19 patients.
- Compared to severely ill COVID-19 patients and to patients with fatal outcomes, the (non-vaccinated) asymptomatic COVID-19 patients presented more functional cross-reactive CD4^+^ and CD8^+^ T cells targeting conserved epitopes from structural, non-structural, and regulatory SARS-CoV-2 protein antigens.
- The cross-reactive SARS-CoV-2 epitopes that recalled the strongest CD4^+^ and CD8^+^ T cell responses in unexposed healthy donors (HD) were the most strongly associated with better disease outcomes seen in asymptomatic COVID-19 patients.
- Compared to severely ill COVID-19 patients and to patients with fatal outcomes, the (non-vaccinated) asymptomatic COVID-19 patients presented higher rates of co-infection with the α-CCCs strains.
- Compared to patients with mild or asymptomatic COVID-19, severely ill symptomatic patients and patients with fatal outcomes had more exhausted SARS-CoV-2-speccific CD4^+^ and CD8^+^ T cells that preferentially target cross-reactive epitopes that share high identity and similarity with the β-CCCs strains.

## INTRODUCTION

Coronaviruses (CoVs) are a large family of respiratory viruses that have been circulating for thousands of years and infect a broad range of species including amphibians, birds, and mammals (1, 2). These viruses are enveloped positive-sense, single-stranded RNA viruses with large genomes (26-32 kb) (1, 2). Within the subfamily *Coronavirinae* are four genera, the alpha (α)-, beta (β)-, gamma (*γ*)-, and delta (δ)-coronaviruses (1, 2). Numerous strains of α- and β-CoVs have been isolated from bats that serve as a large (and highly mobile) CoVs reservoir(2). In humans, many α-CoVs and β-CoVs cause a variety of symptoms from a mild cough to severe respiratory diseases(3). Two α-CoV (HCoV-229E and HCoV-NL63) and two β-CoV (HCoV-HKU1 and HCoV-OC43) strains, known as the common cold coronaviruses (CCCs), cause mild upper respiratory symptoms that are usually associated with only mild disease (1, 4).

Until 2002, β-CoVs caused minor medical concerns to humans(1). However, an outbreak of severe acute respiratory syndrome (SARS) with a severe clinical course emerged in China in 2002 and was caused by a novel pathogenic β-CoV, named SARS-CoV-1 or SARS-CoV(1). SARS-CoV-1 spread to nine countries and led to over 8,000 cases and 775 deaths within one year (∼ 9% case fatality rate)(1). A different highly pathogenic zoonotic β-CoV named the Middle East Respiratory Syndrome CoV (MERS-CoV) later emerged from Saudi Arabia in 2012, transmitted to humans through contact with infected dromedary camels, and later led to human-to-human transmission within healthcare settings(1). Within one year, MERS-CoV infected 2,499 individuals and caused over 858 deaths (∼34% case fatality rate). More recently, the highly infectious and pathogenic β-CoV, SARS-CoV-2 that causes COVID-19, emerged in December 2019 (from China), and, as of January 2022, has infected over 290 million individuals and caused more than 5.5 million deaths, with over 825,000 deaths in the United States alone (∼2% case fatality rate).

Mutations and deletions often occur in the genome of SARS-CoV-2, (predominantly in the Spike protein) resulting in more transmissible and pathogenic “variants of concern” (VOCs). Over the last 25 months, twenty SARS-CoV-2 VOCs have been reported around the world. The latest VOC dubbed “Omicron”, with about 50 genetic mutations and a whopping 36 of them in the Spike protein, emerged from South Africa in November 2021. Omicron is less pathogenic, but highly transmissible, and has since led to record-breaking numbers of infections through escape of antibody-mediated immunity elicited by Spike-based vaccines(5).

Within the first two weeks of infection, ∼20% of unvaccinated SARS-CoV-2-positive tested patients become symptomatic and develop severe COVID-19 disease(6). Symptoms begin with a mild upper respiratory syndrome and may develop into severe respiratory distress and death, especially in immunocompromised individuals and those with pre-existing co-morbidities. While the mechanisms that lead to severe COVID-19 remain to be fully elucidated, immune dysregulations are associated with the pathogenesis of COVID-19 including: (*i*) virus-specific adaptive immune responses that can trigger pathological processes characterized by localized or systemic inflammatory processes(7); (*ii*) increased levels of pro-inflammatory cytokines (8-10); and (*iii*) a broad lymphopenia (11-16). Nevertheless, the role of CD4^+^ and CD8^+^ T cells in COVID-19 disease remains controversial (15-32). CD4^+^ and CD8^+^ T cells specific to SARS-CoV-2 have been reported to be associated with less severe symptoms (33-42). Conversely, SARS-CoV-2-specific CD4^+^ and CD8^+^ T cells have been attributed to poor COVID-19 disease outcomes (43-48). We and others have recently detected cross-reactive SARS-CoV-2-specific CD4^+^ and CD8^+^ T cells directed toward epitopes that are conserved between human CoVs and animal SARS-Like Coronaviruses (SL-CoVs), not only from COVID-19 patients, but also from a significant proportion of healthy individuals that have never been exposed to SARS-CoV-2 infection (1, 36, 37, 49-55). However, whether these pre-existing cross-reactive CD4^+^ and CD8^+^ T cells in healthy individuals and COVID-19 patients play a role in disease protection or pathogenicity caused by SARS-CoV-2 infection has not been elucidated.

In the present study, we characterized the frequency, phenotype, and function of cross-reactive SARS-CoV-2-specific CD4^+^ and CD8^+^ T cells, targeting a large set of SARS-CoV-2 genome-wide conserved epitopes, in a cohort of 147 non-vaccinated COVID-19 patients that were divided into six groups, based on disease severity. Our results showed that, compared to asymptomatic COVID-19 patients, the critically ill COVID-19 patients and those who died from COVID-19 complications: (*i*) had a broad lymphopenia and lowest frequencies of cross-reactive SARS-CoV-2-specific CD4^+^ and CD8^+^ T cells; (*ii*) presented the highest frequencies of SARS-CoV-2-specific CD4^+^ and CD8^+^ T cells with phenotypic and functional exhaustion; (*iii*) appeared to have fewer co-infections with α-CCCs strains. In contrast, compared to severely ill COVID-19 patients and patients with fatal outcomes, the (non-vaccinated) asymptomatic COVID-19 patients: (*i*) presented higher rates of co-infection with the α-CCCs strains; and (*ii*) developed more functional SARS-CoV-2-specific CD4^+^ and CD8^+^ T cells preferentially targeting cross-reactive epitopes that were the most highly recognized by T cells from healthy donors.

## MATERIALS & METHODS

### Human study population cohort and HLA genotyping

From July 2020 to November 2021, we enrolled 600 patients at the University of California Irvine Medical Center with mild to severe COVID-19 symptoms for this study. All subjects were enrolled under an approved Institutional Review Board–approved protocol (IRB#-2020-5779). A written informed consent was obtained from participants prior to inclusion. SARS-CoV-2 positivity was defined by a positive RT-PCR on a respiratory tract sample. In this study, none of the patients received any COVID-19 vaccine. Patients for which the given amount of blood was insufficient (i.e., less than 6ml) were removed. Of the remaining individuals, 147 were genotyped for HLA-A*02:01^+^ or/and HLA-DRB1*01:01^+^ (**Supplemental Fig. S1**). The average days between the report of their first symptoms and the blood sample drawing was 4.8 days (**Table 1** and **Supplemental Table 1**). Following patient discharge, they were divided into groups depending on the severity of their symptoms and their intensive care unit (ICU) and intubation (mechanical ventilation) status (**Table 1** and **Supplemental Table 1**). Scoring was performed by the medical practitioners at the hospital. Accordingly, 9 individuals were asymptomatic (ASYMP – severity score 0) while 138 were symptomatic (SYMP). Among these 138 patients, 12 patients developed very-mild COVID-19 – severity score 1 –, 64 patients developed mild/moderate disease – severity score 2 –, and 62 patients had severe or very severe symptoms. We subsequently divided these severely-ill patients into three groups: patients in ICU – severity score 3, 21 patients – patients in ICU who required mechanical ventilation – severity score 4, 15 patients –, and patients who later died from COVID-19 – severity score 5, 26 patients. Across all the 147 patients 28% were white Hispanic, 22% Hispanic Latino, 16% were Asian, 13% white Caucasian, 8% were mixed Afro-American and Hispanic, 5% were Afro-American, 2% were mixed Afro-American and Caucasian, 1% were of Native Hawaiian and Other Pacific Islander descent and 6% of the patients categorized their race/ethnicity as Other. Forty-one percent were females, and 59% were males with an age range of 19-91 (median 56 and average 55). Compared to the asymptomatic group, the symptomatic patients (i.e., mild, moderate and severe/very severe) were on average older (median: 57 vs. 27) and had a higher percentage of comorbidities (2.4 comorbidities on average vs. 0.7 for the ASYMP), including diabetes (50% to 0%), obesity (46% to 44%), hypertension (39% to 11%), cardiovascular and coronary-arterial diseases (33% and 28% to 0% and 0%, respectively), kidney, diseases (20% to 0%), asthma (18% to 11%) and cancers (10% to 0%) (**Table 1**). The clinical and demographic characteristics of the COVID-19 patients with respect to age, gender, HLA-A*02:01 or HLA-DRB1*01:01 distribution, disease severity, comorbidity, symptoms and symptoms onset, length of stay in the hospital, pulmonary function, immunological parameters, and blood components are summarized in **Table 1** and **Supplemental Table 1**. Fifteen liquid-nitrogen frozen PBMCs samples (blood collected pre-COVID-19 in 2018) from HLA-A*02:01^+^/HLA-DRB1*01:01^+^ unexposed healthy individuals (Healthy Donor: HD – 8 males, 7 females; median age: 54 (20-76)) were used as controls to measure recalled SARS-CoV-2 cross-reactive T cell responses.

**Table 1:**
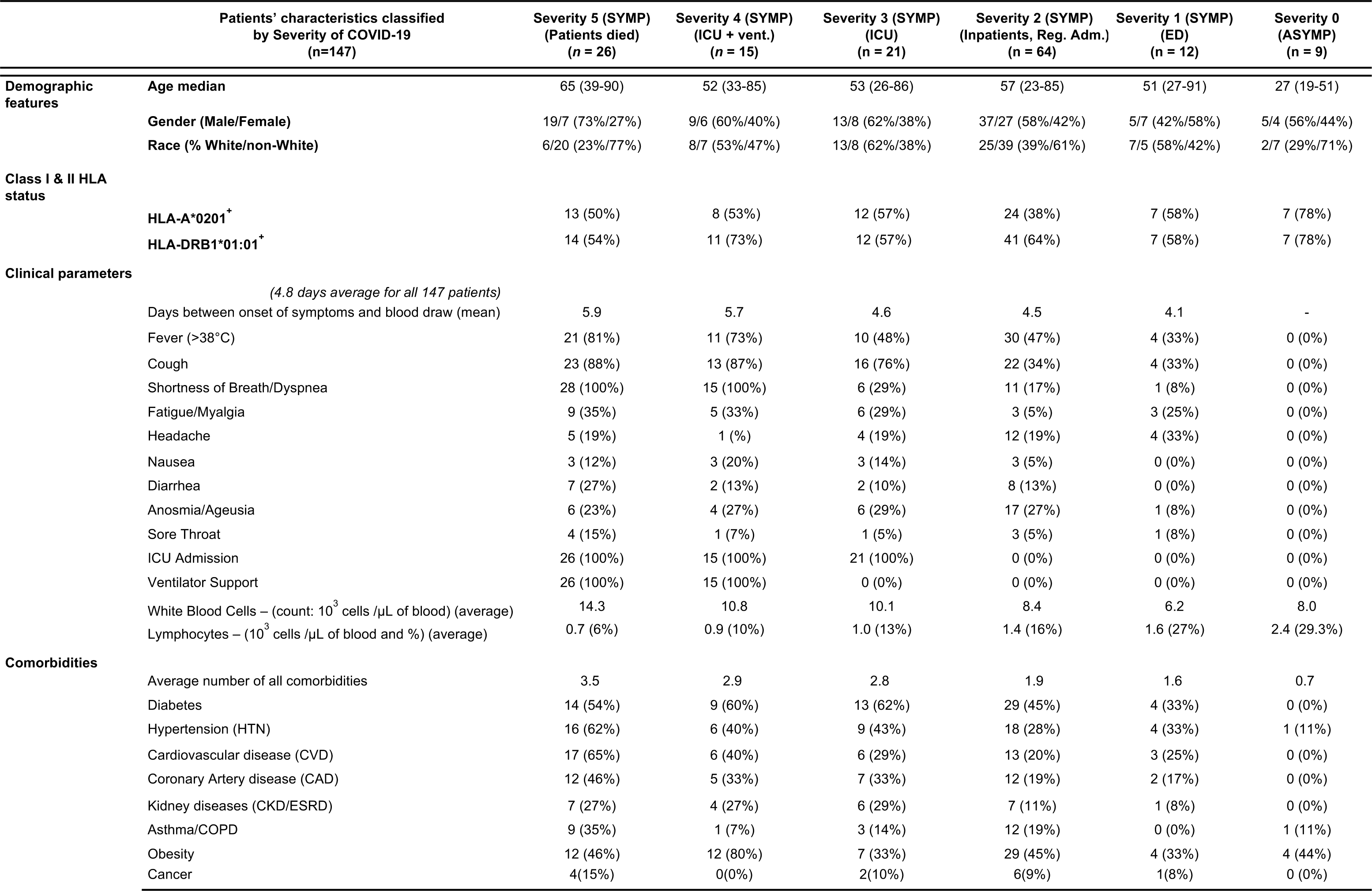
Demographic features, age, HLA-genotyping, clinical parameters, onset of symptoms and prevalence of comorbidities in COVID-19 patients enrolled in the study: Once discharged, patients were scored (“severity score”, or “category of COVID-19 severity”) on a scale from 0 to 5 according to the apparition of symptoms, their hospital department attribution and if they went under Intensive Care Unit (ICU – severity 3), needed life support i.e., mechanical ventilation (at any point during their stay – severity 4) or died from COVID-19 (severity 5). Patients who had no symptoms (severity score 0) were also called ASYMP (asymptomatic) whereas all the patients who developed symptoms (independently of the disease severity) were broadly categorized as SYMP. For SYMP patients who did not go to the ICU, we had: ED = patients who went to the Emergency Department, got screened COVID-19 but did not stay in the hospital for regular admission (severity 1). Reg. Adm. = patients who were admitted for Regular Admission to stay in the hospital to treat their COVID-19 but did not go to ICU (severity 2). Except for the age, the onset of symptoms, the WBCs and lymphocytes count and the total number of comorbidities, all the parameters displayed in the table (demographic features, HLA-genotyping, clinical parameters, and prevalence of comorbidities) represent the number of patients within each category of disease severity and the percentages in brackets (rebased to the total number of patients in the corresponding category). For the age parameter, median values are shown for each category of disease severity along with ranges (between brackets). Per category: time between the onset of symptoms and the blood draw are day-average numbers; the WBCs & lymphocytes counts are averages per µL of blood; and the total number of comorbidities is the average of the sums of each patient’s comorbidities.

The class-II HLA status of each patient was first screened for HLA-DRB1*01:01 by PCR (**Supplemental Fig. S1A**) with the protocol described by(56), using sense primer 5’-TTGTGGCAGCTTAAGTTTGAAT-3’ and two antisense primers: 5’-ACTGTGAAGCTCTCACCAAC-3’ (“*primer 3a*”) and 5’-GGCCCGCCTCTGCTCCA-3’ (“*primer 3c*”). For class-I HLA, the screening was first performed (two-digit level) by HLA-A*02 flow staining (*data not shown*, mAbs clone BB7.2, BioLegend, San Diego, CA). The four-digit class-I HLA-A*02:01 subtype was subsequently screened by PCR (**Supplemental Fig. S1B**) on blood samples from the subjects using the PCR method described previously(57) (sense primer 5′-CCTCGTCCCCAGGCTCT-3′ and antisense 5′-TGGCCCCTGGTACCCGT-3′).

### T cell epitopes screening, selection and peptide synthesis

Peptide-epitopes from twelve SARS-CoV-2 proteins, including 27 9-mer long CD8^+^ T cell epitopes (ORF1ab_84-92_, ORF1ab_1675-1683_, ORF1ab_2210-2218_, ORF1ab_2363-2371_, ORF1ab_3013-3021_, ORF1ab_3183-3191_, ORF1ab_3732-3740_, ORF1ab_4283-4291_, ORF1ab_5470-5478_, ORF1ab_6419-6427_, ORF1ab_6749-6757_, S_2-10_, S_691-699_, S_958-966_, S_976-984_, S_1000-1008_, S_1220-1228_, E_20-28_, E_26-34_, M_52-60_, M_89-97_, ORF_63-11_, ORF7b_26-34_, ORF8a_31-39_, ORF8a_73-81_, ORF_103-11_ and ORF_105-13_) and 16 13-mer long CD4^+^ T cell epitopes (ORF1a_1350-1365_, ORF1a_1801-1815_, ORF1ab_5019-5033_, ORF1ab_6088-6102_, ORF1ab_6420-6434_, S_1-13_, E_20-34_, E_26-40_, M_176-190_, ORF_612-26_, ORF7a_1-15_, ORF7a_3-17_, ORF7a_98-112_, ORF7b_8-22_, ORF8b_1-15_ and N_388-403_) that we formerly identified were selected as described previously(1). Briefly, we first identified consensus protein sequences after performing a sequence conservation analysis between SARS-CoV-2– and SARS-&-MERS-Like-CoVs–protein sequences obtained from human, bat, pangolin, civet, and camel(1). Subsequently, we used multiple databases and algorithms (SYFPEITHI, MHC-I / MHC-II Binding Predictions, Class I Immunogenicity, Tepitool, TEPITOPEpan and NetMHC) to screen conserved CD8^+^ T cell candidate epitopes predicted to bind the 5 most frequent HLA-A class I alleles (HLA-A*01:01, HLA-A*02:01, HLA-A*03:01, HLA-A*11:01, HLA-A*23:01) and conserved CD4^+^ T cell candidate epitopes predicted to bind 5 class II alleles with large population coverage regardless of race and ethnicity, namely DRB1*01:01, HLA-DRB1*11:01, HLA-DRB1*15:01, HLA-DRB1*03:01, HLA-DRB1*04:01(1). The Epitope Conservancy Analysis tool was used to compute the degree of identity of CD8^+^ T cell and CD4^+^ T cell epitopes within a given protein sequence of SARS-CoV-2 set at 100% identity level(1). Peptides were synthesized as previously described(1) (21^st^ Century Biochemicals, Inc, Marlborough, MA). The purity of peptides determined by both reversed-phase high-performance liquid chromatography and mass spectroscopy was over 95%. Peptides were first diluted in DMSO and later in PBS (1 mg/mL concentration).

### Blood Differential Test (BDT)

Total White Blood Cells (WBCs) count and Lymphocytes count per µL of blood were performed by the clinicians at the University of California Irvine Medical Center, using CellaVision^TM^ DM96 automated microscope. Monolayer smears were prepared from anticoagulated blood and stained using the May Grunwald Giemsa (MGG) technique. Subsequently, slides were loaded onto the DM96 magazines and scanned using a 10-x objective focused on nucleated cells to record their exact position. Images were obtained using the 100-x oil objective and analyzed by Artificial Neural Network (ANN).

### Peripheral blood mononuclear cells isolation and T cell stimulation

Peripheral blood mononuclear cells (PBMCs) from COVID-19 patients were isolated from the blood using Ficoll (GE Healthcare) density gradient media and transferred into 96 well plates at a concentration of 2.5 × 10^6^ viable cells per ml in 200µl (0.5 × 10^6^ cells per well) of RPMI-1640 media (Hyclone) supplemented with 10% (v/v) FBS (HyClone), Sodium Pyruvate (Lonza), L-Glutamine, Nonessential Amino Acids, and antibiotics (Corning). A fraction of the blood was kept separated to perform HLA genotyping of the patients and select only the HLA-A*02:01 and/or DRB1*01:01 positive individuals (**Supplemental Fig. S1**). Subsequently, cells were then stimulated with 10µg/ml of each one of the 43 individual T cell peptide-epitopes (27 CD8^+^ T cell peptides and 16 CD4^+^ T cell peptides) and incubated in humidified 5% CO_2_ at 37°C (**Supplemental Fig. S2A**). Incubation times were for either 72 hours straight prior to the cells being stained by flow cytometry analysis, or for 24 hours before being transferred in IFN-γ ELISpot plates for an additional 48 hours (i.e., for a total of 72 hours stimulation in both conditions). Same isolation protocol was followed for HD samples taken in 2018. Ficoll were kept in frozen in liquid nitrogen in FBS DMSO 10%; after thawing, HD PBMCs were stimulated in the same manner for IFN-*γ* ELISpot.

### ELISpot assay

COVID-19 patients were first screened for their HLA status: out of 147 samples, 92 were DRB1*01:01^+^ and 71 were HLA-A*02:01^+^ whereas, 16 patients were screened positive for both (**Supplemental Fig. S1** and **Supplemental Table 1**). The ninety-two DRB1*01:01 positive individuals were used to assess the CD4^+^ T-cell response against our SL-CoVs-conserved SARS-CoV-2-derived class-II restricted epitopes by IFN-*γ* ELISpot. Seven individuals were asymptomatic – severity score 0; 7 patients had very-mild COVID-19 – severity score 1; 41 showed moderate disease – severity score 2; and 37 patients developed severe or very severe symptoms, divided into 3 groups: 12 patients in ICU without mechanical ventilation – severity score 3; 11 patients in ICU who required mechanical ventilation – severity score 4; and 14 patients who died from COVID-19 – severity score 5.

Similarly, we assessed the CD8^+^ T-cell response against our SL-CoVs-conserved SARS-CoV-2-derived class-I restricted epitopes in the seventy-one HLA-A*02:01 positive individuals. Seven individuals were asymptomatic – severity score 0; 7 patients had very-mild COVID-19 – severity score 1; 24 showed moderate disease – severity score 2; and 33 patients developed severe or very severe symptoms, divided into 3 groups: 12 patients in ICU without mechanical ventilation – severity score 3; 8 patients in ICU who required mechanical ventilation – severity score 4; and 13 patients who later died from COVID-19 – severity score 5.

All ELISpot reagents were filtered through a 0.22 µm filter. Wells of 96-well Multiscreen HTS Plates (Millipore, Billerica, MA) were pre-wet with 30% ethanol for 60 seconds and then coated with 100 µl primary anti-IFN-*γ* antibody solution (10 µg/ml of 1-D1K coating antibody from Mabtech, Cincinnati, OH) OVN at 4°C. After washing, the plate was blocked with 200 µl of RPMI media plus 10% (v/v) FBS for 2 hours at room temperature to prevent nonspecific binding. Following the blockade, twenty-four hours peptide-stimulated cells from the patients PBMCs (0.5 x 10^6^ cells/well) were transferred into the ELISpot coated plates. PHA stimulated or non-stimulated cells (DMSO) was used as positive or negative controls of T cells activation, respectively. After incubation in a humidified chamber with 5% CO2 at 37°C for an additional 48 hours, cells were washed using PBS and PBS-Tween 0.02% solution. Next, 100 µl of biotinylated secondary anti-IFN-*γ* antibody (1 µg/ml, clone 7-B6-1, Mabtech) in blocking buffer (PBS 0.5% FBS) were added to each well. Following a 2-hour incubation and washing, wells were incubated with 100 µl of HRP-conjugated streptavidin (1:1000) for 1 hour at room temperature. Lastly, wells were incubated for 15-30 minutes at room temperature with 100 µl of TMB detection reagent and spots were counted both manually and by an automated ELISpot reader counter (ImmunoSpot Reader, Cellular Technology, Shaker Heights, OH).

### Flow cytometry analysis

After 72 hours of stimulation with each individual SARS-CoV-2 class-I or class-II restricted peptide, PBMCs from the same 147 patients were stained for surface markers detection (0.5 x 10^6^ cells) and subsequently analyzed by flow cytometry (**Supplemental Fig. S2**). First, the cells were stained with a live/dead fixable dye (Zombie Red dye, 1/800 dilution – BioLegend, San Diego, CA) for 20 minutes at room temperature, to exclude dying/apoptotic cells. Cells were then stained for 45 minutes at room temperature with five different HLA-A*02*01 restricted tetramers and/or five HLA-DRB1*01:01 restricted tetramers (PE-labelled) specific toward the SARS-CoV-2 CD8^+^ T cell epitopes Orf1ab_2210-2218_, Orf1ab_4283-4291_, S_976-984_, S_1220-1228_, ORF_103-11_ and toward the CD4^+^ T cell epitopes ORF1a_1350-1365_, S_1-13_, E_26-40_, M_176-190_, ORF_612-26_, respectively. Cells were alternatively stained with the EBV BMLF-1_280–288_-specific tetramer(58) for controls. We optimized our tetramer staining according to instructions published by Dolton et al.(59). Subsequently, we used the following anti-human antibodies for surface-marker staining: anti-CD45 (BV785, clone HI30 – BioLegend), anti-CD3 (Alexa700, clone OKT3 – BioLegend), anti-CD4 (BUV395, clone SK3 – BD), anti-CD8 (BV510, clone SK1 – BioLegend), anti-TIGIT (PercP-Cy5.5, clone A15153G – BioLegend), anti-TIM-3 (BV 711, clone F38-2E2 – BioLegend), anti-PD1 (PE-Cy7, clone EH12.1 – BD), anti-CTLA-4 (APC, clone BNI3 – BioLegend), anti-CD138 (APC-Cy-7, clone 4B4-1 – BioLegend) and anti-CD134 (BV650, clone ACT35 – BD). mAbs against these various cell markers were added to the cells in phosphate-buffered saline (PBS) containing 1% FBS and 0.1% sodium azide (fluorescence-activated cell sorter [FACS] buffer) and left for 30 minutes at 4°C. At the end of the incubation period, the cells were washed twice with FACS buffer and fixed with paraformaldehyde 4% (PFA, Affymetrix, Santa Clara, CA). A total of ∼200,000 lymphocyte gated PBMCs (140,000 alive CD45^+^) were acquired by Fortessa X20 (Becton Dickinson, Mountain View, CA) and the subsequent analysis performed using FlowJo software (TreeStar, Ashland, OR). Gating strategy is detailed in **Supplemental Fig. S2B**.

### TaqMan quantitative polymerase reaction assay for the detection of common-cold in COVID-19 patients

To detect common-cold coronaviruses co-infections in COVID-19 patients, Taqman PCR assays were performed on a total of 85 patients distributed into each different category of disease severity (9 ASYMP, 6 patients of category 1, 32 patients of category 2, 9 patients of category 3, 15 patients of category 4 and 14 patients of category 5). Nucleic acid was first extracted from each blood sample using QIAamp MinElute Virus Spin kits (Qiagen, Mississauga, Ontario, Canada) according to the manufacturer’s instructions. Subsequently, extracted RNA samples were quantified using the Qubit and BioAnalyzer. cDNA was synthesized from 10 μL of RNA eluate using random hexamer primers and SuperScript II Reverse Transcriptase (Applied Biosystems, Waltham, MA). The subsequent RT–PCR screening of the enrolled subjects for the four CCCs was performed using the following specific sets of primers and probes: for **CCC-229E**: forward primer 5’-CAGTCAAATGGGCTGATGCA-3’, reverse primer 5’-AAAGGGCTATAAAGAGAATAAGGTATTCT-3’ and Taq-Man probe 5’-NED-CCCTGACGACCACGTTGTGGTTCA-MGBNFQ-3’; for CCC-OC43: forward primer 5’-CGATGAGGCTATTCCGACTAGGT-3’, reverse primer 5’-CCTTCCTGAGCCTTCAATATAGTAACC-3’ and Taq-Man probe FAM-TCCGCCTGGCACGGTACTCCCT-MGBNFQ-3’; for CCC-NL63: forward primer 5’-ACGTACTTCTATTATGAAGCATGATATTAA-3’, reverse primer 5’-AGCAGATCTAATGTTATACTTAAAACTACG-3’ and Taq-Man probe 5’-NED-ATTGCCAAGGCTCCTAAACGTACAGGTGTT-MGBNFQ-3’; and finally for CCC-HKU1: forward primer 5’-CCATTACAAGCCATAAGAGAACAAAC-3’, reverse primer 5’-TATGTGTGGCGGTTGCTATTATGT-3’ and Taq-Man probe 5’-FAM-TTGCATCACCACTGCTAGTACCACCAGG-TAMRA-3’)(60).

CCC-229E, CCC-OC43, and CCC-NL63 RT-PCR assays were performed using the following conditions: 50°C for 15 minutes followed by denaturation at 95°C for 2 minutes, 40 cycles of PCR performed at 95°C for 8 seconds, extending and collecting fluorescence signal at 60°C for 34 seconds(61). For CCC-HKU1, the amplification conditions were 48°C for 15 minutes, followed by 40 cycles of 94°C for 15 seconds and 60°C for 15 seconds. For each virus, when the Ct-value generated was less than 35, the specimen was considered positive. When the Ct-value was relatively high (35 ≤ Ct < 40), the specimen was retested twice and considered positive if the Ct-value of any retest was less than 35(62).

### SARS-CoV-2 epitope identity analysis with the corresponding best-matching CCCs-peptides from each of the four CCCs and peptide similarity score calculation

To assess the % identity (%id) of our SL-CoVs-conserved SARS-CoV-2-derived CD4^+^ and CD8^+^ T cell peptide-epitopes, we first identified the best matching CCCs peptide across the CCCs proteomes. The full CCCs proteomes sequences were obtained from the National Center for Biotechnology Information (NCBI) GenBank (with the following accession authentication numbers: MH940245.1 for CCC-HUK1, MN306053.1 for CCC-OC43, KX179500.1 for CCC-NL63 and MN306046.1 for CCC-229E. We processed this in three steps. (**1**) Corresponding CCCs peptides were determined after proteins sequences alignments of all four homologous CCCs proteins plus the SARS-CoV-2 related one using various Multiple Sequences Alignments (MSA) algorithms ran in JALVIEW, MEGA11 and M-coffee software’s (i.e. ClustalO, Kalign3 and M-coffee – the latter computing alignments by combining a collection of Multiple Alignments from a Library constituted with the following algorithms: T-Coffee, PCMA, MAFFT, ClustalW, Dialigntx, POA, MUSCLE, and Probcons). In addition, we confirmed our results with global and local Pairwise alignments (Needle and Water algorithms ran in Biopython). In the case of obtaining different results with the various algorithms, the epitope sequence with the highest BLOSUM62-sum score compared to the SARS-CoV-2 epitope set as reference was selected (**Supplemental Table 5**). We calculated % of identity and similarity score **S^s^** with its related SARS-CoV-2 epitope, for each of these CCCs peptides (**Supplemental Table 5**). The peptide similarity score S^s^ calculation is based on Sune Frankild et al. method(63) and the BLOSUM62 matrix to calculate a BLOSUM62 sum (using the *Bio.SubsMat.MatrixInfo* package in Biopython) between a pair of peptides (peptide “x” from SARS-CoV-2 and “y” from one CCC) and compare their similarity. 0 ≤ S^s^ ≤ 1: the closest S^s^ is to 1, the highest is the potential for T cell cross-reactivity response toward the related pair of peptide(63). We used a threshold of S^s^≥0.8 to discriminate between highly similar and non-similar peptides. (**2**) Then, we examined if other parts of each CCCs proteome (without restricting our search only to peptides present in CCCs homologous proteins) could contain better matching peptides than the CCCs peptides reported in **Supplemental Table 5** (found after MSA). First, for each one of our 16 CD4^+^ and 27 CD8^+^ SARS-CoV-2 epitopes, we spanned the entire proteome of each CCCs using the Epitope Conservancy Tool (ECT: http://tools.iedb.org/conservancy/ – with a conservancy threshold of 20%). All the CCCs peptides from the top query (i.e., with the highest % of identity) were reported for each four CCCs in the **Supplemental Table 6**. Second, among these returned top queries (peptides with the same highest % of identity), we picked the one with the highest similarity score S^s^ (bolded in **Supplemental Table 6** – *right column*). (**3**) We compared this peptide with the one previously found in **Supplemental Table 5** based on MSA. When both methods returned the same peptide (from the same protein), we kept it (peptides highlighted in beige in **Supplemental Table 6**, reported in **Supplemental Table 3**). When both matching peptides (using the two different methods) were found to be different, we compared (***i***) %id_MSA_ with %id_ECT_ then (***ii***) S^s^_MSA_ with S^s^_ECT_. If %id_MSA_ ≤ %id_ECT_ but S^s^_MSA_ ≥ S^s^_ECT_, we kept the CCCs peptide found following the MSA method; however, if %id_MSA_ ≤ %id_ECT_ and S^s^_MSA_ < S^s^_ECT_, we then picked the CCC peptide found using the ECT instead of the one found using MSA (peptides not highlighted in **Supplemental Table 3**).

Using the %id and the calculated similarity score with the SARS-CoV-2 epitopes, all related CCCs best matching peptides were reported in **Supplemental Table 3.** They were then evaluated based on their potential of inducing a cross-reactive T cell response, as shown in **Supplemental Table 4**: (**0**): CCC best matching peptide with low to no potential to induce a cross-reactive response toward the corresponding SARS-CoV-2 epitope and vice-versa (%id with the corresponding SARS-CoV-2 epitope < 67% AND similarity score S^s^ < 0.8); (**0.5**): CCC best matching peptide that may induce a cross-reactive response (%id with the corresponding SARS-CoV-2 epitope ≥ 67% OR similarity score S^s^ ≥ 0.8); (**1**): CCC best matching peptide very likely to induce a cross-reactive response (%id ≥ 67% AND S^s^ ≥ 0.8).

### Identification of potential cross-reactive peptide in common human pathogens and vaccines

We took advantage of the database generated by Pedro A. Reche(64). Queries to find matching peptides with our SARS-CoV-2-derived CD4^+^ and CD8^+^ epitopes were performed from the data gathered; only peptides sharing a %id ≥ 67% with our corresponding SARS-CoV-2 epitope were selected (**Supplemental Table 7)**. The corresponding similarity score S^s^ was calculated, and results reported in **Supplemental Table 4**.

### Statistical analyses

To assess the potential linear negative relationship between COVID-19 severity and the magnitude of each SARS-CoV-2 epitope-specific T cell response, correlation analysis using GraphPad Prism version 8 (La Jolla, CA) were performed to calculate the Pearson correlation coefficients (R), the coefficient of determination (R^2^) and the associated *P*-value (correlation statistically significant for *P* ≤ 0.05). The slope (S) of the best-fitted line (dotted line) was calculated in Prism by linear-regression analysis. Same statistical analysis was performed to compare the cross-reactive pre-existing T cell response in unexposed HD with the slope S (magnitude of correlation between this epitope-specific T cell response in SARS-CoV-2 infected patients and the protection against severe COVID-19). Absolute WBC and lymphocytes cell numbers (per µL of blood, measured through BDT), corresponding lymphocytes percentages/ratio, Flow Cytometry data measuring CD3^+^/CD8^+^/CD4^+^ cell percentages and the percentages detailing the magnitude (Tetramer^+^ T cell %) and the quality (% of PD1^+^/TIGIT^+^, CTLA-4^+^/TIM3^+^ or AIMs^+^ cells) of the CD4^+^ and CD8^+^ SARS-CoV-2 specific T cells, were compared across groups and categories of disease severity by one-way ANOVA multiple tests. ELISpot SFCs data were compared by Student’s *t*-tests. Data are expressed as the mean ± SD. Results were considered statistically significant at *P* ≤ 0.05. To evaluate whether the differences in frequencies of RT-PCR positivity to the four CCCs across categories of disease severity was significant, we used the Chi-squared test (when comparing three groups of COVID-19 severity) or the Fisher’s exact test (when comparing two groups of COVID-19 severity).

## RESULTS

### 1. Lower magnitudes of SARS-CoV-2-specific CD4^+^ T cell responses detected in severely ill COVID-19 patients compared to mild and asymptomatic COVID-19 patients

We first compared SARS-CoV-2-specific CD4^+^ T cell responses in symptomatic vs. asymptomatic COVID-19 patients (**Fig. 1**). We used 16 recently identified HLA-DR-restricted CD4^+^ T cell epitopes that are highly conserved between human SARS-CoVs and animal SL-CoVs(1). We enrolled 92 non-vaccinated HLA-DRB1*01:01^+^ COVID-19 patients, genotyped using PCR (**Supplemental Fig. S1**), and divided into six groups, based on the level of severity of their disease (from severity 5 to severity 0, assessed at discharge – **Table 1**). Severity 5: patients who died from COVID-19 complications; Severity 4: infected COVID-19 patients with severe disease that were admitted to the intensive care unit (ICU) and required ventilation support; Severity 3: infected COVID-19 patients with severe disease that required enrollment in ICU, but without ventilation support; Severity 2: infected COVID-19 patients with moderate symptoms that involved a regular hospital admission; Severity 1: infected COVID-19 patients with mild symptoms; and Severity 0: infected individuals with no symptoms. Detailed clinical, gender and demographic characteristics of this cohort of COVID-19 patients are shown in **Table 1** and **Supplemental Table 1**. Fresh peripheral blood mononuclear cells (PBMCs) were isolated from these COVID-19 patients, on average within 4.8 days after reporting their first symptoms (**Table 1**). PBMCs were then stimulated *in vitro* for 72 hours using each of the 16 CD4^+^ T cell epitopes, as detailed in *Materials* & *Methods* and in **Supplemental Fig. S2**. Subsequently, we determined the numbers of responding IFN-*γ*-producing CD4^+^ T cells, induced in each of the six groups, by each of the 16 HLA-DR-restricted epitopes (quantified in ELISpot assay as the number of IFN-*γ*-spot forming cells, or “SFCs”) (**Fig. 1**).

**Figure 1.**
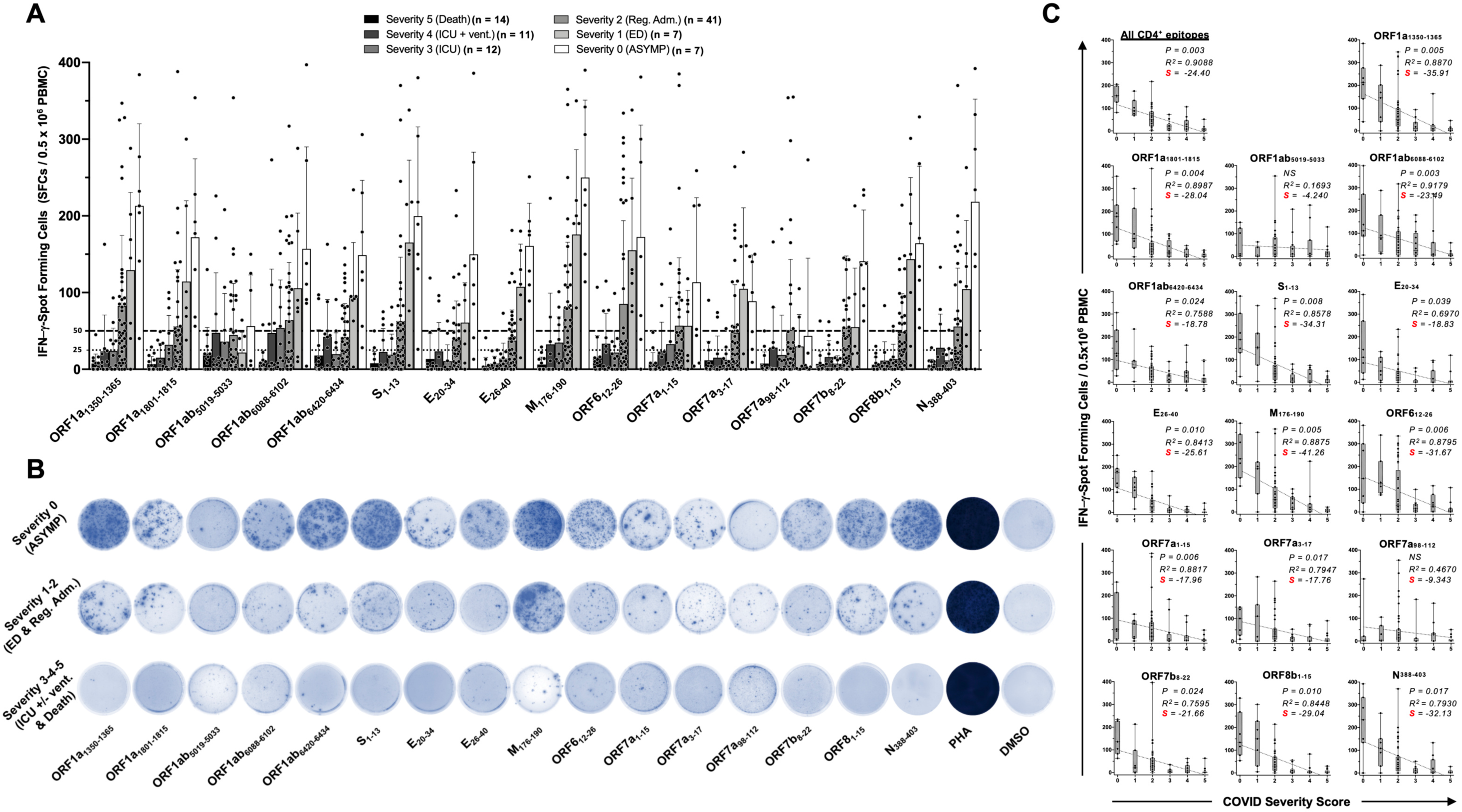
Magnitude of the IFN-γ CD4^+^ T cell responses specific to 16 conserved SARS-CoV-2-derived epitopes in COVID-19 patients with various degrees of disease severity: PBMCs from COVID-19 patients HLA-DRB1*01:01 positives (*n* = 92) were isolated and stimulated for a total of 72 hours with 10µg/ml of each of the 16 class-II restricted peptides, previously identified as SARS-CoV-2-derived CD4^+^ T cell epitopes (see the experimental design in **Supplemental Fig. S2A**). (**A**, **B** and **C**) The number of IFN-*γ*-producing cells were quantified using ELISpot assay for each one of the 92 patients, assigned into one of the six categories of disease severity (scored 0 to 5 –incrementing with the severity): panel (**A**) shows the average/mean numbers (± SD) of IFN-*γ*-spot forming cells (SFCs) after CD4^+^ T cell peptide-stimulation in COVID-19 patients who ended up with various levels of disease severity (for each epitope, categories 0 to 5 being identified by six columns on a grayscale: from black -severity 5, to white -severity 0). Dotted lines represent an arbitrary threshold set to evaluate the relative magnitude of the response: a mean SFCs between 25 and 50 correspond to a medium/intermediate response, whereas a strong response is defined for a mean SFCs > 50 per 0.5 x 10^6^ stimulated PBMCs. (**B**) Representative spots images of the IFN-*γ* response following PBMCs peptide-stimulation from three patients, each one falling into one of the following three groups of disease category: ASYMP patients (severity score 0), patients who developed mild to moderate COVID-19 disease (severity score 1 and 2) and patients who developed severe to very severe disease (severity scores 3 to 5). PHA was used as positive control of T-cell activation. SFCs from the negative control (DMSO – no peptide stimulation) were subtracted from the SFCs counts of peptides-stimulated cells. In chart (**C**), each graph named for each peptide/epitope-stimulation represent the correlations between the overall number of the SARS-CoV-2-specific IFN-*γ*-producing CD4^+^ T cells and the corresponding COVID-19 disease severity. For all graphs are indicated: the coefficient of determination (R^2^) calculated from the Pearson correlation coefficients (R – showed in **Supplemental Table 2**), its associated *P*-value and the slope (S) of the best-fitted line (dotted line) calculated by linear-regression analysis. The gray-hatched boxes in the correlation graphs extend from the 25th to 75th percentiles (hinges of the plots) with the median represented as a horizontal line in each box and the extremity of the vertical bars showing the minimum and maximum values. Results were considered statistically significant at *P* ≤ 0.05.

Overall, the highest frequencies of IFN-*γ*-producing CD4^+^ T cells (determined as mean SFCs > 50 per 0.5 x 10^6^ PBMCs fixed as threshold) were detected early in COVID-19 patients with less severe disease (i.e., severity 0, 1 and 2, **Figs. 1A** and **1B**). In contrast, the lowest frequencies of IFN-*γ*-producing CD4^+^ T cells directed toward SARS-CoV-2 epitopes were detected in the remaining two groups of severely ill symptomatic COVID-19 patients (i.e., severity 3 and 4, mean SFCs < 50) and the group of patients with fatal outcomes (i.e., severity 5, mean SFCs < 25).

To determine a potential linear correlation between the magnitude of CD4^+^ T cell responses directed toward each of the 16 highly conserved SARS-CoV-2 epitopes and COVID-19 disease severity, we performed a Pearson correlation analysis, where a negative correlation is usually considered strong when the coefficient R is comprised between -0.7 and -1(65). Except for the ORF1ab_5019-5033_ and ORF7a_98-112_ epitopes, we found a strong negative linear correlation between the magnitude of IFN-*γ*-producing CD4^+^ T cells against all the remaining 14 epitopes and the severity of COVID-19 disease (**Fig. 1C**). Consequently, a positive correlation existed between the magnitude of CD4^+^ T cell responses specific to 14 CD4^+^ T cell epitopes and the “natural protection” seen in asymptomatic COVID-19 patients. This correlation existed regardless of whether CD4^+^ T cells target structural or non-structural SARS-CoV-2 antigens. However, both the Pearson correlation coefficient (R) (**Supplemental Table 2**) and the coefficient of determination (R^2^, **Fig. 1C**) give a measure of linearity of a possible two-way linear association but do not quantify the “magnitude” of this relationship, which is given by the slope (S) of the best-fitted line (linear regression) shown in **Fig. 1C**. For any T cell epitope-specific response where a negative correlation with the onset of severe symptoms is significant, a strongly negative slope S indicates that the higher the initial T cell response against this epitope, the lower the associated COVID-19 disease severity score. **Supplemental Table 2** illustrates in SARS-CoV-2-infected patients, the epitope-specific CD4^+^ T cell responses that were the most negatively associated with subsequent severe COVID-19 (using a blue/red color code).

An early IFN-*γ*-producing CD4^+^ T cell response specific to M_176-190_, ORF1a_1350-1365_, S_1-13_, N_388-403_, ORF_612-26_, and to a slightly lesser extent to ORF8b_1-15_, and ORF1a_1801-1815_, were associated with a low COVID-19 severity score (i.e., negatively correlated with a R close to -1) and a very strong negative slope (-41.26 < S < -28.04). Comparatively, the CD4^+^ T cell responses against E_26-40_, ORF1ab_6088-6102_, ORF7b_8-22_, E_20-34_, ORF1ab_6420-6434_, ORF7a_1-15_ and ORF7a_3-17_, were also negatively associated with severe disease in patients, but to a lesser degree (relatively less negative slope: -25.61 < S < -17.76) (**Fig 1C** and **Supplemental Table 2**). In contrast, no significant correlation was found between the magnitude of IFN-*γ*-producing CD4^+^ T cell responses directed towards ORF1ab_5019-5033_ and ORF7a_98-112_ epitopes and the disease severity (*P* > 0.05). For the ORF1ab_5019-5033_ and ORF7a_98-112_ epitopes, where the slope was comparatively weak: only slightly negative with S > -10 (**Fig. 1C** and **Supplemental Table 2**).

Taken together, these results demonstrate an overall lower magnitude of SARS-CoV-2-specific CD4^+^ T cell responses in symptomatic and severely ill COVID-19 patients. In contrast, higher magnitudes of SARS-CoV-2-specific CD4^+^ T cell responses were detected in asymptomatic COVID-19 patients. The findings suggest an important role of SARS-CoV-2-specific CD4^+^ T cells directed against both structural and non-structural antigen in protection from severe COVID-19 symptoms and highlights the importance of rapidly mounting strong CD4^+^ T cell responses directed towards SARS-CoV-2 epitopes that are highly conserved between human SARS-CoVs and animal SL-CoVs.

### 2. Lower magnitudes of SARS-CoV-2-specific CD8^+^ T cell responses detected in severely ill COVID-19 patients compared to mild and asymptomatic COVID-19 patients

We next compared SARS-CoV-2-specific CD8^+^ T cell responses in symptomatic vs. asymptomatic COVID-19 patients (**Fig. 2**). We used 27 recently identified HLA-A*0201-restricted CD8^+^ T cell epitopes that are highly conserved between human SARS-CoVs and animal SL-CoVs(1). We enrolled 71 non-vaccinated HLA-A*0201^+^ COVID-19 patients, genotyped using PCR (**Supplemental Fig. S1**), and divided into 6 groups based on disease severity, as stated above (i.e., severity 5 to severity 0, **Table 1** and **Supplemental Table 1**). Fresh PBMCs, isolated from COVID patients on average 4.8 days after reporting their first symptoms, were stimulated *in vitro* for 72 hours using each of the 27 CD8^+^ T cell epitopes, as described in *Materials* & *Methods* (**Supplemental Fig. S2**). The numbers of responding IFN-*γ*-producing CD8^+^ T cells, induced in each of the six groups, by each of the 27 HLA-A*0201-restricted epitopes were determined by ELISpot, as previously detailed (**Fig. 2**).

**Figure 2:**
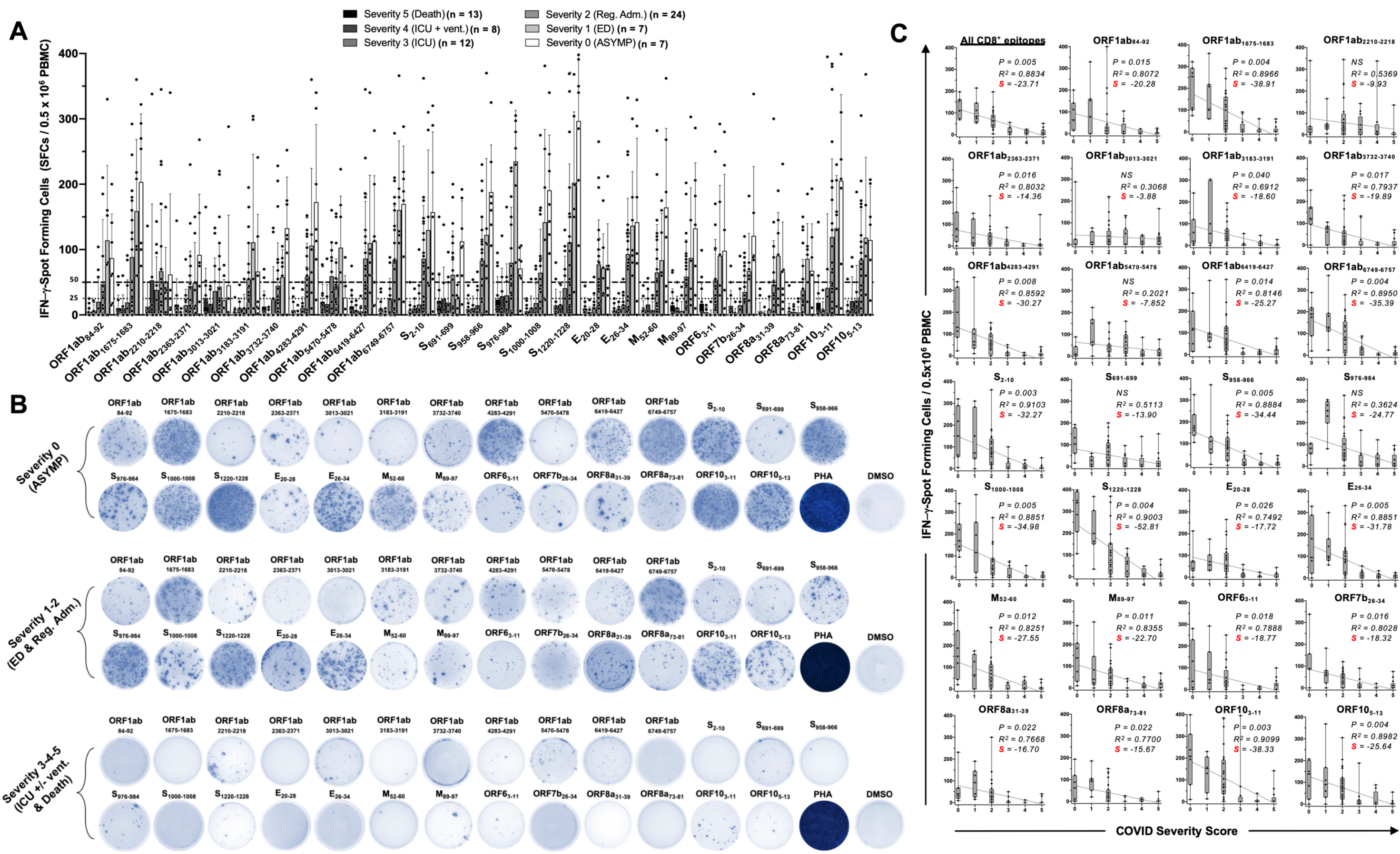
Magnitude of the IFN-γ CD8^+^ T cell responses specific to specific to 27 conserved SARS-CoV-2-derived epitopes in COVID-19 patients with various degrees of disease severity: PBMCs from COVID-19 patients HLA-A*02:01 positives (*n* = 71) were isolated and stimulated for a total of 72 hours with 10µg/ml of each of the 27 class-I restricted peptides, previously identified as SARS-CoV-2-derived CD8^+^ T cell epitopes (**Supplemental Fig. S2A**). (**A, B and C**) The number of IFN-*γ*-producing CD8^+^ T cells were quantified using ELISpot assay for each one of the 71 patients and for each disease severity category: panel (**A**) shows the average/mean numbers (± SD) of IFN-*γ*-spot forming cells (SFCs) after CD8^+^ T cell peptide-stimulation in COVID-19 patients who ended up with various levels of disease severity (using the same legend than before). Dotted lines represent arbitrary threshold set to evaluate the relative magnitude of the response: a mean SFCs between 25 and 50 correspond to a medium/intermediate response whereas a strong response is defined for a mean SFCs > 50 per 0.5 x 10^6^ stimulated PBMCs. (**B**) Representative spots images of the IFN-*γ* response following PBMCs peptide-stimulation from three patients, each one falling into one of the following three groups of disease category already described in the first figure. PHA is used as positive control of T-cell activation and SFCs from the negative control (DMSO – no peptide stimulation) were subtracted from the SFCs counts of peptides-stimulated cells. Chart (**C**) shows the correlation graphs for each peptide/epitope linking the overall number of the corresponding SARS-CoV-2-specific IFN-*γ*-producing CD8^+^ T cells with the disease severity. For all graphs are indicated: the coefficient of determination (R^2^) calculated from the Pearson correlation coefficients (showed in **Supplemental Table 2**), its associated *P*-value and the slope (S) of the best-fitted line (dotted line) calculated by linear-regression analysis. The gray-hatched boxes in the correlation graphs extend from the 25th to 75th percentiles (hinges of the plots) with the median represented as a horizontal line in each box and the extremity of the vertical bars showing the minimum and maximum values. Results were considered statistically significant at *P* ≤ 0.05.

Overall, highest frequencies of functional IFN-*γ*-producing CD8^+^ T cells (mean SFCs > 50 per 0.5 x 10^6^ PBMCs) were detected early in the three groups of COVID-19 patients with no to low severity disease (i.e., severity 0, 1 and 2, **Figs. 2A** and **2B**). In contrast, the lowest frequencies of functional IFN-*γ*-producing CD8^+^ T cells were detected in the 2 groups of severely ill symptomatic COVID-19 patients (i.e., severity 3 and 4, mean SFCs < 50) and in patients with fatal outcomes (i.e., severity 5, mean SFCs < 25). These results suggest that, like CD4^+^ T cells, there was an association between low magnitudes of SARS-CoV-2-specific CD8^+^ T cell responses and severe COVID-19 disease onset. Moreover, there was an association between high magnitudes of SARS-CoV-2-specific CD8^+^ T cell responses and a no to low COVID-19 severity of disease. This association was regardless of whether CD8^+^ T cells targeted epitopes from structural, non-structural, or regulatory SARS-CoV-2 protein antigens.

Out of the 27 CD8^+^ T cell epitopes, there was a significant negative linear correlation between CD8^+^ T cell responses specific to 22 epitopes and COVID-19 disease severity (**Figs. 2A** and **2B**). For these 22 epitopes, the Pearson correlation coefficients (R) ranged from -0.8314 to -0.9541 and slopes (S) of the best-fitted lines comprised between -14.36 and -52.81 (**Supplemental Table 2**). For the remaining 5 epitopes (ORF1ab_2210-2218_, ORF1ab_3013-3021_, ORF1ab_5470-5478_, S_691-699_, and S_976-984_), no significant linear correlation was observed. Nonetheless, among these 5 epitopes, the slope for ORF1ab2210-2218, ORF1ab3013-3021 and ORF1ab5470-5478 was comparatively less negative (S > -10) (**Fig. 2C** and **Supplemental Table 2**). Also, although we could not establish any significant linear correlation relationship between CD8^+^ T cell responses against S_691-699_ or S_976-984_ and disease severity, more-complex (non-linear) associations might exist. For example, the magnitude of the S_976-984_-specific IFN-*γ*-producing CD8^+^ T cell response followed a clear downside trend as the disease severity increased in severely ill symptomatic COVID-19 patients and patients with fatal outcomes (i.e., severity 3, 4 and 5) (**Fig. 2A** and **Fig. 2C**: S_S976-984_ = -24.77).

Taken together, these results demonstrate that in COVID-19 patients, low SARS-CoV-2-specific CD8^+^ T cell responses were more commonly associated with severe disease onset. In contrast, higher magnitudes of SARS-CoV-2-specific CD8^+^ T cell responses were detected in asymptomatic COVID-19 patients. These findings suggest that, in addition to CD4^+^ T cells, SARS-CoV-2-specific CD8^+^ T cells directed against both structural and non-structural antigens play an important role in protection from COVID-19 severe symptoms and highlights the importance of rapidly mounting strong CD8^+^ T cell responses directed towards SARS-CoV-2 epitopes that are highly conserved between human SARS-CoVs and animal SL-CoVs.

### 3. A broad lymphopenia and low frequencies of SARS-CoV-2 specific CD4^+^ and CD8^+^ T cells associated with severe disease in COVID-19 patients

We next sought to determine whether the low magnitude of SARS-CoV-2 specific IFN-*γ*-producing CD4^+^ and CD8^+^ T cell responses detected in severely ill and fatal COVID-19 patients was a result of an overall deficit of CD4^+^ and CD8^+^ T cells. Using a blood differential test (BDT), we compared the absolute numbers of white blood cells (WBCs) and blood-derived lymphocytes, *ex vivo,* in the six groups of COVID-19 patients (i.e., severity 0, 1, 2, 3, 4, and 5, **Fig. 3A**).

**Figure 3:**
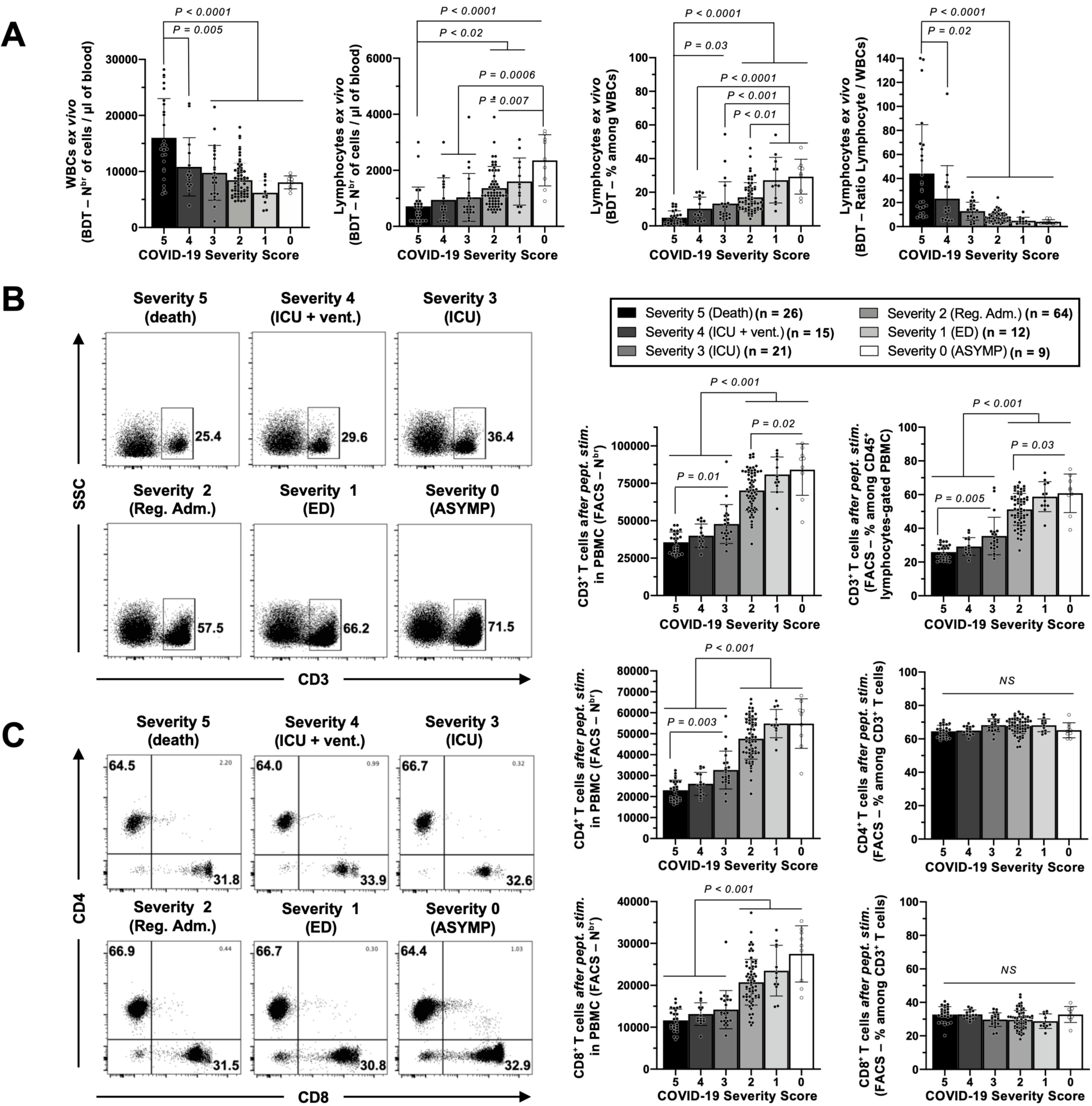
Frequencies and absolute numbers of white blood cells, lymphocytes and CD3^+^/CD4^+^/CD8^+^ T cells in the blood of COVID-19 patients with various degrees of disease severity: (**A**) Numbers of white blood cells (WBCs) and total lymphocytes per µl of blood (*two left panels*) and percentages and ratios of total lymphocytes among WBCs (*two right panels*) measured ex-vivo by blood differential test (BDT) in COVID-19 patients (*n* = 147) who ended up with various severity of disease. Details of BDT results for each patient are shown in Supplemental Table 1. (**B and C**) Percentages and numbers of total CD3^+^ T cells (**B**), CD4^+^ and CD8^+^ T cells (**C**) measured by flow cytometry from COVID-19 patients’ PBMCs with various severity scores after 72 hours SARS-CoV-2 specific peptide-stimulation with our pool of 16 CD4^+^ and 27 CD8^+^ peptides. For both **B** and **C** i.e., respectively for the CD3^+^ staining gated from the CD45^+^ parental population and for the CD4^+^/CD8^+^ staining gated from the CD3^+^ parental population (detailed gating strategy is shown in Supplemental Fig. S2B), *right panels* show representatives dot plots from patients with disease severity scores from 0 (ASYMP) to 5 (patients who died from COVID-19). *Left panels* show associated columns graphs with averages/means of numbers and frequencies (from the gated parental populations) of the CD3^+^ T cells (**B**) and the CD4^+^/CD8^+^ T cells (**C**). Data are expressed as the mean ± SD. Results were considered statistically significant at *P* ≤ 0.05 (one-way ANOVA).

Blood samples were isolated from COVID-19 patients on average 4.8 days after reporting their first symptoms. A significant increase (between ∼1.5- and ∼2.6-fold) in the numbers of WBCs was detected in patients with fatal outcomes, (i.e., severity 5) when compared with all the 4 other groups of COVID-19 patients (i.e., severity 0, 1, 2, 3 and 4; *P* ≤ 0.02, **Fig. 3A** –*left panel*). In addition, we found a significantly lower absolute numbers of total lymphocytes circulating in the blood of patients with fatal outcomes compared to patients with mild disease (severity 1 and 2: ∼1.9- and ∼2.3-fold decrease – *P* < 0.02) or to asymptomatic patients (severity 0: ∼3.3-fold decrease – *P*<0.0001) (**Fig. 3A** – *second panel from left*). Comprehensively, the most severely ill patients (severity 3, 4 and 5) had significantly fewer blood-derived lymphocytes than patients developing little to no disease (severity 0, 1 and 2). As a result, the more severe the disease, the lower the percentage of blood-derived lymphocytes among WBCs (**Fig. 3A** – *third panel from left*) and the higher the ratio of lymphocyte/WBCs (**Fig. 3A** – *fourth panel from left*). Overall, these results indicate that severely ill COVID-19 patients and COVID-19 patients with fatal outcomes not only have a general and abrupt blood leukocytosis but also lymphopenia, as early as 4.8 days after reporting their first symptoms.

Moreover, using flow cytometry (gating shown **Supplemental Fig. S2**), we found a CD3^+^ T cell lymphopenia in severely ill COVID-19 patients, which was positively associated with the onset of severe disease (**Fig. 3B**). On average, the three groups of severely ill COVID-19 patients and COVID-19 patients with fatal outcomes (Severity 3, 4 and 5) had a ∼1.9-fold decrease in both frequency and absolute number of CD3^+^ T cells compared to COVID-19 patients with low to no severe disease (Severity 0, 1 and 2, **Fig. 3B**, *P* < 0.001). A similar trend was observed for the numbers of both CD4^+^ and CD8^+^ T cells (**Fig. 3C** – *left column graph*). However, no significant difference was detected across groups of disease severity in the percentages of both CD4^+^ and CD8^+^ among CD3^+^-gated T cells (*P* > 0.05, **Fig. 3C** – *right column graph*), demonstrating that both the CD4^+^ and CD8^+^ T cells were similarly reduced early on in patients who developed severe COVID-19.

Finally, we determined the frequencies of SARS-CoV-2 tetramer-positive T cells specific to 5 different CD4^+^ and 5 different CD8^+^ SARS-CoV-2-derived SL-CoVs-conserved epitopes (**Fig. 4**) after 72 hours of corresponding peptide-stimulation (**Supplemental Fig. S2**). Respectively: ORF1a_1350-1365_, S_1-13_, E_26-40_, M_176-190_ and ORF_612-26_ for the DRB1*01:01-restricted CD4^+^ epitopes (**Fig. 4A**) and Orf1ab_2210-2218_, Orf1ab_4283-4291_, S_976-984_, S_1220-1228_ and ORF_103-11_ for the A*02:01-restricted CD8^+^ epitopes (**Fig. 4B**). In the most severely ill patients and patients with fatal outcomes (severity 3, 4 and 5), we found a significant decrease in the frequencies of tetramer-positive CD4^+^ T cells specific to all the 5 SARS-CoV-2 DRB1*01:01-restricted epitopes compared to patients with mild disease (severity 1, 2 – *P* ≤ 0.01) or no disease (severity 0 – *P* ≤ 0.002) (**Fig. 4A**). Similarly, we found a significant decrease in the frequencies of tetramer-positive CD8^+^ T cells specific to 3 out of the 5 SARS-CoV-2 A*02:01-restricted epitopes (Orf1ab_4283-4291_, S_1220-1228_ and ORF_103-11_ – **Fig. 4B**) in the most severely ill symptomatic patients (severity 3, 4 and 5) compared to patients with mild disease (severity 1,2 – *P* ≤ 0.03) or asymptomatic patients (*P* < 0.001). Except for ORF1ab_2210-2219_ and S_976-984_ epitopes (for which we showed in **Fig. 2** that there was no significant negative correlation between the associated IFN-*γ* response and disease severity), the lowest frequencies of epitope-specific CD4^+^ and CD8^+^ T cells were detected in the group of severely ill symptomatic COVID-19 and in patients with fatal outcomes (i.e., severity 3, 4 and 5; **Fig. 4**). This contrasts with similar frequencies of EBV BMLF-1280– 288-specific CD8^+^ T cells detected across the groups of COVID-19 patients regardless of disease severity, indicating that the decrease in the frequencies among T cells in severely ill COVID-19 patients particularly affected T cells specific to SARS-CoV-2 epitopes (**Supplemental Fig. S3A**). Finally, similar to the reduction of IFN-*γ*-producing SARS-CoV-2-specific CD4^+^ and CD8^+^ T cells seen in severely ill symptomatic COVID-19 and in patients with fatal outcomes (**Figs. 1** and **2** above), the reduction in the frequencies of SARS-CoV-2-specific CD4^+^ and CD8^+^ T cells appeared regardless of whether T cells targeted structural or non-structural antigens (**Figs. 4A** and **4B**).

**Figure 4:**
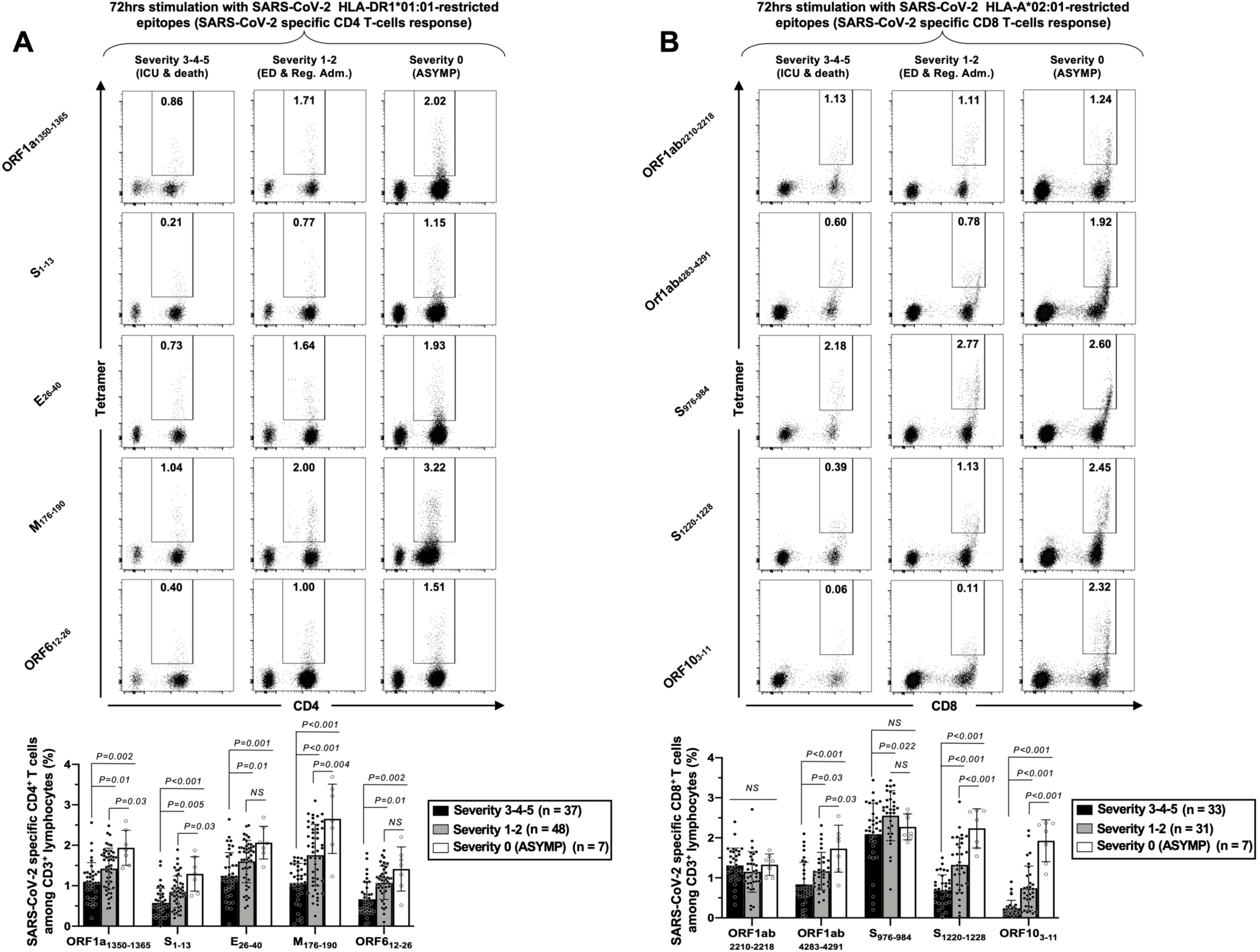
Frequencies of SARS-CoV-2-derived epitopes-specific CD4^+^ and CD8^+^ T cells in COVID-19 patients with various degrees of disease severity: PBMCs from HLA-DRB1*01:01 positive (n=92) (**A**) or HLA-A*02:01 positive (*n* = 71) (**B**) COVID-19 patients divided in three groups of various disease severity scores (severity 0 (ASYMP), severity 1-2 (mild/moderate) and severity 3-4-5 (severe disease)) were isolated and stimulated 72 hours with 10 μg/ml of each one of five SL-CoVs-conserved SARS-CoV-2-derived CD4^+^ peptides/epitopes and each one of five SL-CoVs-conserved SARS-CoV-2-derived CD8^+^ peptides/epitopes listed in the figure. The patients’ PCMCs were stained, analyzed by flow cytometry, and subsequently gated according to the protocol and gating strategy described in Supplemental Fig. S2. The 10 epitopes were chosen among our 16 CD4^+^ and 27 CD8^+^ SL-CoVs-conserved SARS-CoV-2-derived epitopes according to the corresponding tetramer availability. *Upper panel* in (**A**) and *upper panel* in (**B**) shows representative dot plots of the tetramer staining against the five CD4^+^ epitopes and the five CD8^+^ epitopes (respectively) for the three groups of disease severity. *Lower panels* in (**A**) and (**B**) demonstrate associated columns graphs with averages/means of tetramer-positive T cell frequencies. Data are expressed as the mean ± SD. Results were considered statistically significant at *P* ≤ 0.05 (one-way ANOVA).

Taken together, the findings: (*i*) confirmed previous reports demonstrating broad early lymphopenia (and leukocytosis) in severely ill COVID-19 patients (11-15, 66, 67); (*ii*) demonstrated that the decrease of bulk CD3^+^ T cell lymphocytes numbers (affecting both CD4^+^ and CD8^+^ T cells equally) in severely ill COVID-19 patients was one major cause of this lymphopenia, but more importantly; (*iii*) that SARS-CoV-2-specific CD4^+^ and CD8^+^ T cells responding to conserved epitopes from structural, non-structural and regulatory protein antigens were even more reduced (particularly decreased relative to total reduction in T cells) in severely ill patients and in COVID-19 patients with fatal outcomes, this soon after reporting their first symptoms.

### 4. Compared with asymptomatic COVID-19 patients, severely ill symptomatic COVID-19 patients have higher frequencies of phenotypically and functionally exhausted SARS-CoV-2-specific CD4^+^ and CD8^+^ T cells

We next determined whether the low magnitudes of SARS-CoV-2-specific IFN-*γ*-producing T cell responses (**Figs. 1** and **2**) and low frequencies of tetramer-positive SARS-CoV-2-specific CD4^+^ and CD8^+^ T cells (**Fig. 4**) detected in severely ill symptomatic COVID-19 and in patients with fatal outcomes could be the result of phenotypic and functional exhaustion of SARS-CoV-2-specific CD4^+^ and CD8^+^ T cells. Using flow cytometry, we determined the co-expression of four main exhaustion receptors (PD-1, TIM3, TIGIT and CTLA4) and two activation markers (AIMs) CD138 (4-1BB) and CD134 (OX40) on tetramer-positive CD4^+^ T cells specific to five structural and non-structural SARS-CoV-2 epitopes (ORF1a_1350-1365_, S_1-13_, E_26-40_, M_176-190_ and ORF_612-26_, **Fig. 5**) and on tetramer-positive CD8^+^ T cells specific to five structural and non-structural SARS-CoV-2 epitopes (Orf1ab_2210-2218_, Orf1ab_4283-4291_, S_976-984_, S_1220-1228_ and ORF_103-11_, **Fig. 6**).

**Figure 5:**
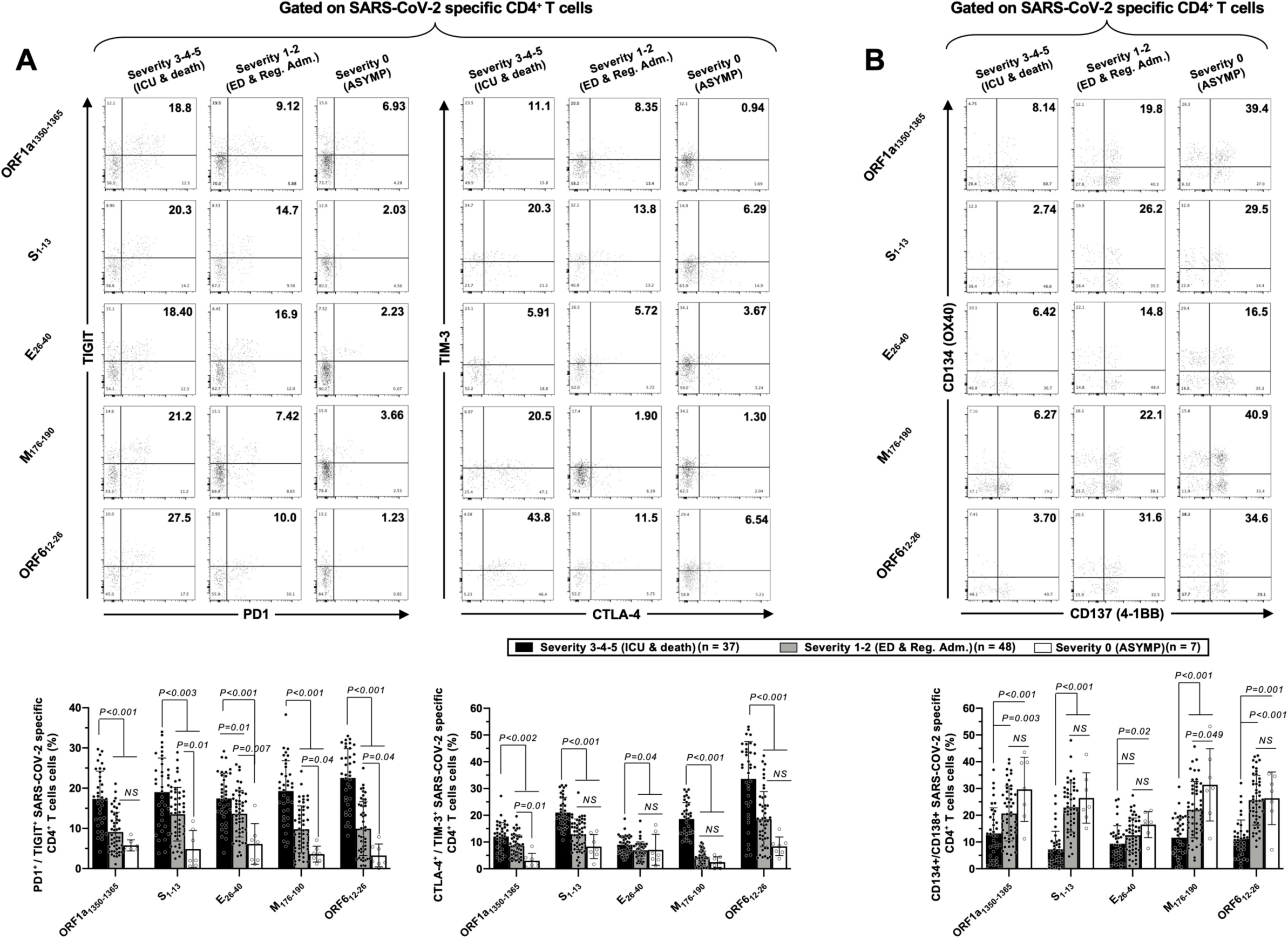
Co-expression of cell surface exhaustion markers PD1, TIGIT, CTLA-4 and TIM-3 by SARS-CoV-2 epitope-specific CD4^+^ T cells in COVID-19 patients with various degrees of disease severity: Experimental design: PBMCs from HLA-DRB1*01:01 positive COVID-19 patients (*n* = 92) divided in three groups of various disease severity scores as in Fig. 4 were isolated and stimulated for 72 hours with 10 μg/ml of each of the 5 SL-CoVs-conserved SARS-CoV-2-derived CD4^+^ T cell epitopes before staining (**Supplemental Fig. S2**) and flow-cytometry acquisition. In (**A**) are shown the frequency of tetramer-specific CD4^+^ cells co-expressing exhaustion receptors PD1 plus TIGIT and TIM-3 plus CTLA-4 after each stimulation, whereas (**B**) shows the frequency of the same cells co-expressing the activation-induced markers (AIMs) CD134 and CD137 after the same treatment. In both (**A** and **B**), *upper panels* depict representative dot plots of the staining and *lower panels* display associated column graphs with averages/means of the frequencies of SARS-CoV-2-specific CD4^+^ T cells co-expressing the exhaustion receptors (in **A**), or the AIMs (in **B**). Data are expressed as the mean ± SD. Results were considered statistically significant at *P* ≤ 0.05 (one-way ANOVA).

**Figure 6:**
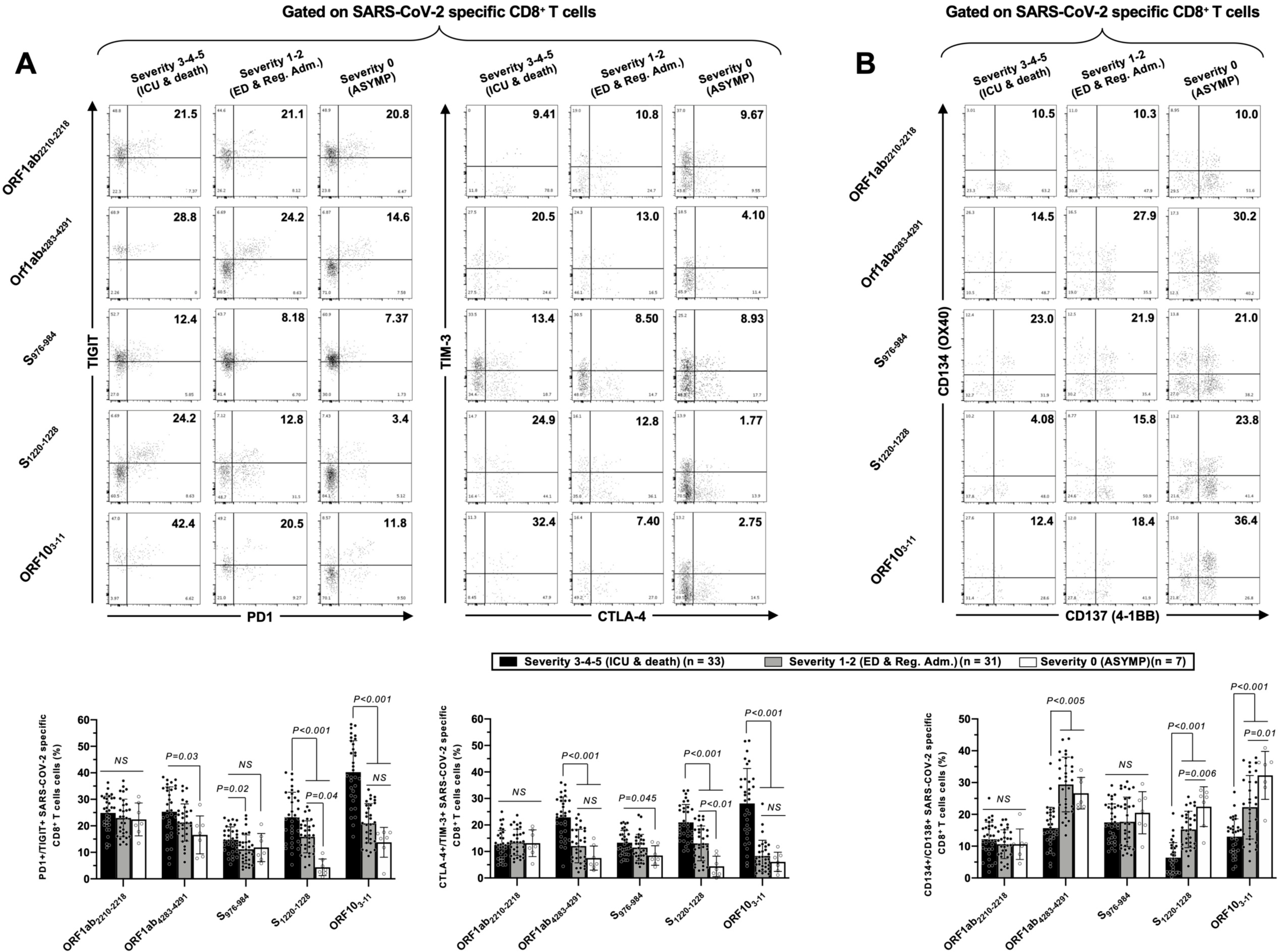
Co-expression of cell surface exhaustion markers PD1, TIGIT, CTLA-4 and TIM-3 by SARS-CoV-2 epitope-specific CD8^+^ T cells in COVID-19 patients with various degrees of disease severity: Experimental design: PBMCs from HLA-A*02:01 positive COVID-19 patients (*n* = 71) divided in three groups of various disease severity scores as in Fig. 4 were isolated and stimulated for 72 hours with 10 μg/ml of each of the 5 SL-CoVs-conserved SARS-CoV-2-derived CD8^+^ T cell epitopes before staining (Supplemental Fig. S2) and flow-cytometry acquisition. In (A) are shown the frequency of tetramer-specific CD8^+^ cells co-expressing exhaustion receptors PD1 plus TIGIT and TIM-3 plus CTLA-4 after each stimulation, whereas (B) shows the frequency of the same cells co-expressing the activation-induced markers (AIMs) CD134 and CD137 after the same treatment. In both (A and B), *upper panels* depict representative dot plots of the staining and *lower panels* display associated columns graphs with averages/means of the frequencies of SARS-CoV-2-specific CD8^+^ T cells co-expressing the exhaustion receptors (in A), or the AIMs (in B). Data are expressed as the mean ± SD. Results were considered statistically significant at *P* ≤ 0.05 (one-way ANOVA).

We detected significantly higher frequencies of phenotypically exhausted SARS-CoV-2-specific CD4^+^ T cells (**Fig. 5A** –up to ∼6.9-fold increase for ORF612-26-specific PD-1^+^TIGIT^+^CD4^+^ T cells and up to ∼7.8-fold increase for M176-190 -specific TIM-3^+^CTLA-4^+^CD4^+^ T cells) in COVID-19 patients with high severity scores (i.e., severity 3, 4 and 5) compared to asymptomatic COVID-19 patients (i.e., severity 0). Similarly, there were significantly higher frequencies of phenotypically exhausted SARS-CoV-2-specific CD8^+^ T cells (**Fig. 6A** –up to ∼3.6-fold increase for S_1220-1228_-specific PD-1^+^TIGIT^+^CD8^+^ T cells and up to ∼4.6-fold increase for S_1220-1228_- and ORF_103-11_-specific TIM-3^+^CTLA-4^+^CD8^+^ T cells) in severely ill COVID-19 and in patients with fatal outcomes compared to asymptomatic COVID-19 patients. Overall, except for Orf1ab_2210-2218_- and S_976-984_-specific-CD8^+^ T cells, the most severely ill patients (severity 3, 4 and 5) had significantly higher frequencies of exhausted T cells co-expressing PD-1^+^TIGIT^+^ or TIM-3^+^CTLA-4 than patients developing little to no disease (severity 0, 1 and 2). That Orf1ab_2210-2218_- and S_976-984_-specific-CD8^+^ T cells showed no significant higher phenotypic exhaustion in severely ill COVID-19 patients was consistent with the observation that CD8^+^ T cell responses to these two epitopes were not associated with severe COVID-19 disease (**Figs. 2** and **4**).

Reflecting the high frequencies of exhausted CD4^+^ and CD8^+^ T cells in severely ill COVID-19 and in patients with fatal outcomes, we also detected the lowest frequencies of functional CD134^+^CD138^+^CD4^+^ T cells (**Fig. 5B**) and CD134^+^CD138^+^CD8^+^ T cells (**Fig. 6B**) in those patients. This applied to CD134^+^CD138^+^CD4^+^ T cells specific to all 5 structural and non-structural SARS-CoV-2 epitopes and for CD134^+^CD138^+^CD8^+^ T cells specific to 3 out of 5 structural and non-structural SARS-CoV-2 epitopes (except Orf1ab_2210-2218_ and S_976-984_-specific CD8^+^ T cells). As expected, there was no difference in phenotypic and functional exhaustion of EBV BMLF-1280–288-specific CD8^+^ T cells across the COVID-19 disease severities (**Supplemental Fig. S3B**), suggesting that the exhaustion in severely ill COVID-19 patients was specific to SARS-CoV-2-specific CD4^+^ and CD8^+^ T cells.

In conclusion, the decrease in the magnitudes of IFN-*γ*-producing SARS-CoV-2 specific T cell responses and in the frequencies of tetramer-positive SARS-CoV-2-specific CD4^+^ and CD8^+^ T cells detected in COVID-19 patients with high severity scores (i.e., severity 3, 4 and 5) was associated with phenotypic and functional exhaustion of CD4^+^ and CD8^+^ T cells specific to those SL-CoVs-conserved epitopes, from both structural and non-structural antigens.

### 5. Compared with asymptomatic COVID-19 patients, severely ill symptomatic COVID-19 patients present lower frequencies of co-infections with α-CCCs

Using RT-PCR, we examined the co-infection with each of the four strains of CCCs (i.e., α-CCC-NL63, α-CCC-229E, β-CCC-HKU1 and β-CCC-OC43) in a cohort of 84 COVID-19 patients divided into six groups with various disease severities (**Fig. 7A**). We found co-infections with α-CCCs strains, to be more common and significantly higher in the asymptomatic COVID-19 patients compared to severely ill COVID-19 patients and in patients with fatal outcomes (**Fig. 7B** *– right panel*: ∼2.6-fold increase in groups 1-2-3 vs. groups 4-5-6 of disease severity; *P* = 0.0418 calculated with Fisher’s exact test). In particular, co-infection with the CoV-229E α-CCC strain was more common and significantly higher in the asymptomatic COVID-19 patients compared to severely ill COVID-19 patients and to patients with fatal outcomes (**Fig. 7C** *– right panels*: ∼4.2-fold increase between asymptomatic and group 4-5-6; *P* = 0.0223 calculated with Chi-squared test). However, there was no significant difference in the frequencies of co-infections with β-CCCs strains (nor with all the four CCC strains altogether) across all severity groups (**Fig. 7B** *– central and left panels* and **Fig. 7C** *– left 2 panels*).

**Figure 7:**
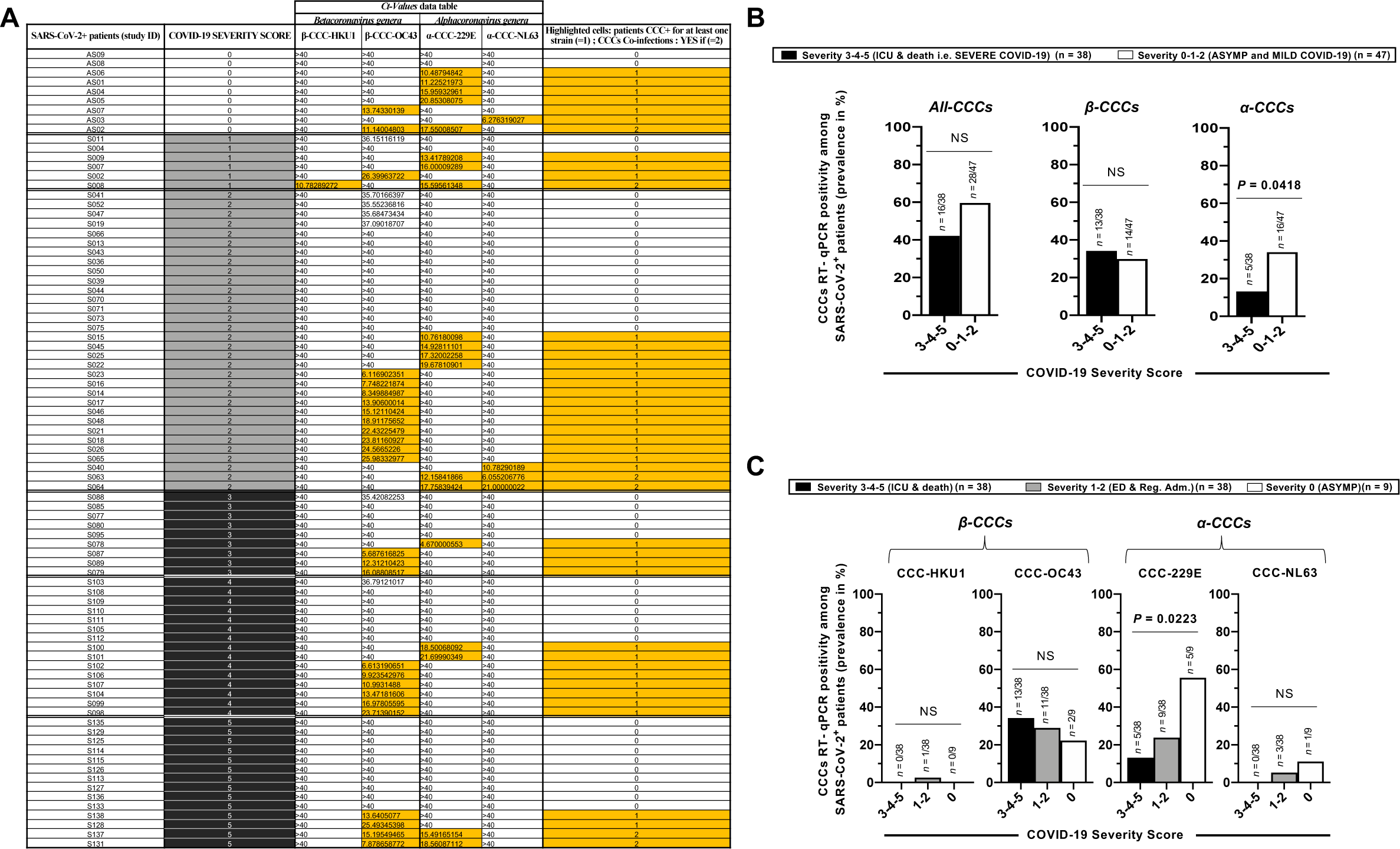
Detection by quantitative RT-PCR of various strains of human common-cold coronaviruses in COVID-19 patients with various degrees of disease severity: For the detection of the four human common-cold coronaviruses (CCC-HKU1, CCC-OC43, CCC-229E and CCC-NL63), COVID-19 patients (n=85) who developed various disease severity were screened through RT–PCR performed after RNA extraction from their blood samples. For each patient (i.e., row), (A) shows the Ct-values generated after RT-PCR amplification. A Ct-value below 35 is synonymous of CCC-positivity for the chosen tested sample/individual (highlighted in light gray in the table). Patients are organized in ascending order of disease severity scores (0 to 5). (B and C) Demonstrate the different genera of common-cold coronaviruses (Beta: CCC-HKU1 and CCC-OC43 – *on the left*; Alpha: CCC-229E and CCC-NL63 – *on the right*) the prevalence (%) of coinfection with these viruses in COVID-19 SARS-CoV-2 infected patients. In (B), patients are divided into 3 groups of disease severity (as in Figs. 4, 5 and 6): severity 0 (ASYMP), severity 1-2 (mild/moderate) and severity 3-4-5 (severe disease). CCCs positivity prevalence are measured for each individual CCC, and *P*-values were calculated using Chi-squared test. In (C), patients are divided into 2 groups of disease severity (severity 0-1-2: ASYMP and mild disease vs. 3-4-5: severe disease) and CCCs positivity prevalence are measured for each CCC genera (Alpha and Beta). *P*-values where here calculated with the Fisher’s exact test. Details of the statistics are provided in the Supplemental Figures. All results were considered statistically significant at *P* ≤ 0.05.

These results indicate that, compared to severely ill COVID-19 patients and to patients with fatal outcomes, the asymptomatic COVID-19 patients presented significantly higher frequencies of co-infections with α-CCCs strains, in general, and with the 229E strain of α-CCCs in particular.

### 6. Compared with severely ill COVID-19 patients, asymptomatic COVID-19 patients develop SARS-CoV-2-specific CD4^+^ and CD8^+^ T cells preferentially targeting CCCs-cross-reactive epitopes that recalled the strongest pre-existing T cells responses in healthy unexposed individuals

We have previously observed that in some unexposed healthy donors (HD), PBMCs stimulation with our CD4^+^ and CD8^+^ SARS-CoV-2-derived epitopes induced an IFN-γ^+^ T cell response(1). We confirmed those results here on 15 additional HD (**Supplemental Fig. S4**). We hypothesized that this pre-existing response predating the COVID-19 pandemic could possibly influence the establishment of the SARS-CoV-2-specific T cell response either positively or negatively, and its effectiveness to prevent the most severe symptoms in infected patients. Therefore, we investigated (**Fig. 8A**) a possible correlation between (***i***) the cross-reactivity of each epitope measured in HD (i.e., the ability of each SARS-CoV-2 CD4^+^ and CD8^+^-derived epitope to recall a SARS-CoV-2 cross-reactive T cell response in unexposed individuals, measured by IFN-γ ELISpot – **Supplemental Fig. S4**) and (***ii***) the percentage of asymptomatic/mild COVID-19 patients (among all asymptomatic/mild COVID-19 patients) for which we could detect a strong IFN-*γ*^+^ CD4^+^ /CD8^+^ T cell response (>50 SFCs – **Figs. 1A** and **2A**), specific to the same epitope. The percentage for each epitope was calculated as follows: number of patients from one category of disease severity with SFCs>50, divided by the total number of patients within this same category.

**Figure 8:**
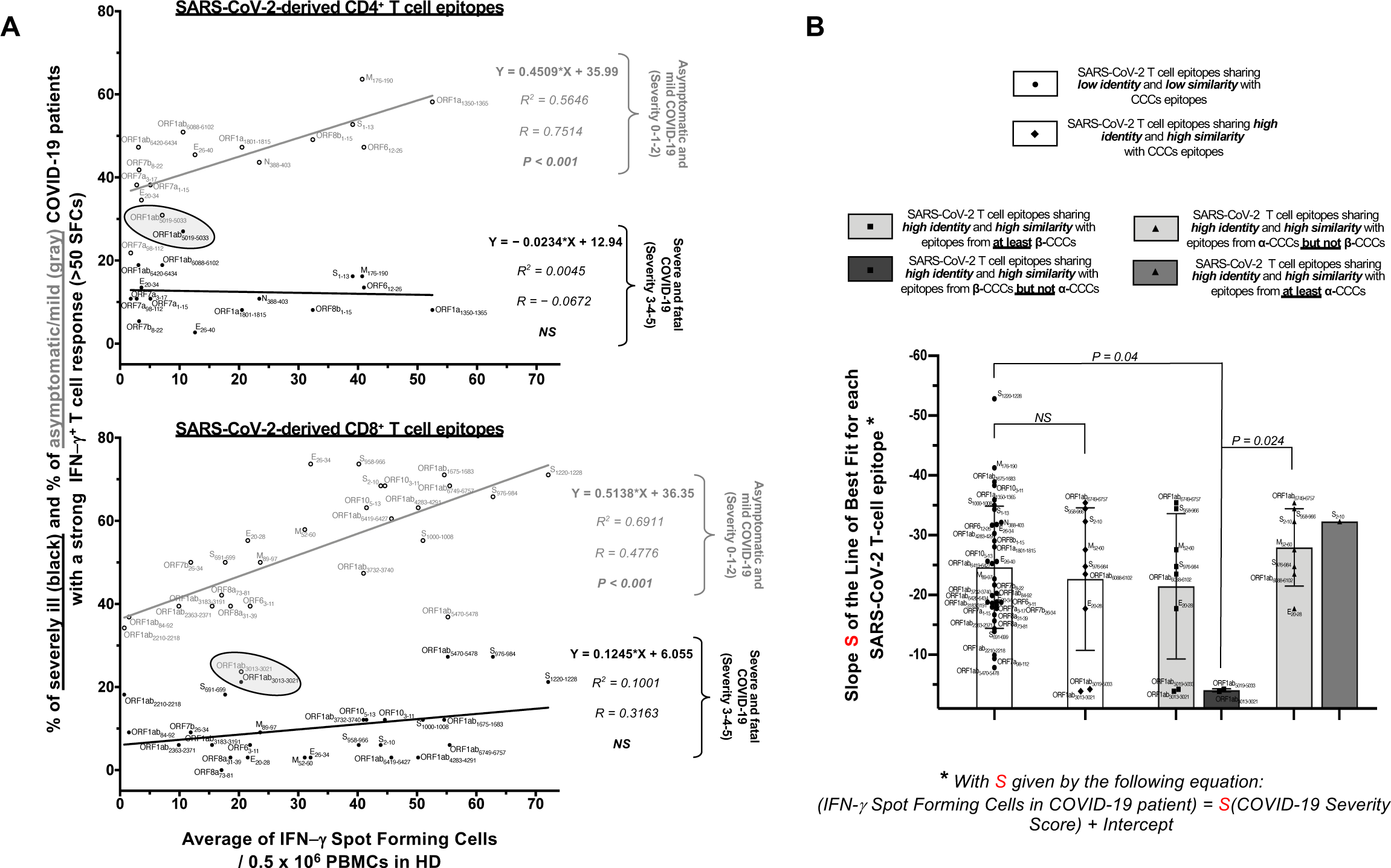
Pre-existing cross-reactive SARS-CoV-2-specific T cell responses in unexposed HD: relations with the T cell responses in SARS-CoV-2 infected patients and the protection against severe COVID-19: (**A**) Both graphs represent the correlations between the cross-reactivity of each epitope in HD (i.e., the average number of SARS-CoV-2-specific cross-reactive IFN-*γ*-producing T cells measured by ELISpot in unexposed healthy donors – Supplemental Fig. S4 – following stimulation with each of the 16 CD4 epitopes, *upper graph*; or the 27 CD8 epitopes, *lower graph*), and the percentage of severely ill (black dots) or asymptomatic/mild (clear dots) COVID-19 patients for which we detected a strong IFN-*γ*^+^ T cell response (>50 SFCs – Figs. 1A and 2A), specific to the corresponding epitope. For both graphs are indicated: the coefficient of determination (R^2^), the Pearson correlation coefficients (R), its associated *P*-value and the equation of the best-fitted line calculated (from linear-regression analysis). (**B**) The slope S of the Line of Best Fit (from Fig. 1C for CD4 epitopes; and from Fig. 2C for CD8 epitopes) is used as a measure of the magnitude of the correlation between the breadth of each epitope-specific T cell response in patients and the corresponding COVID-19 disease severity. For each epitope: the higher S, the more a strong T cell response toward this epitope is associated to a better disease outcome. Here, our SL-CoVs-conserved SARS-CoV-2-derived CD4 and CD8 epitopes are categorized according to their similarity and % of identity with α- and/or β-CCC strains i.e., their cross-reactivity potential to be also recognized by α-CCCs, or β-CCCs, or α-<u>and</u> β-CCCs specific T cells (see **Supplemental Table 4**: high identity is defined for %id>67% and high similarity for a similarity score S^S^≥0.8). Circled epitopes in panel (**A**) are the one found to share high identity and similarity exclusively with β-CCCs (black column). Epitopes’ cross-reactivity categories are compared using one-way ANOVA and results were considered statistically significant at *P* ≤ 0.05.

Within the category of asymptomatic and mild COVID-19 patients, we found statistically significant positive correlations (*P > 0.001*) between the epitope cross-reactivities measured in HD and the percentage of asymptomatic and mild COVID-19 patients that developed a strong IFN-*γ*^+^ T cell response (SFCs>50) specific to SARS-CoV-2-derived CD4^+^ T cell epitopes (**Fig. 8A** *– upper graph, gray line*) or CD8^+^ T cell epitopes (**Fig. 8A** *– lower graph, gray line*). In contrast, no such significant correlations were found within the category of patients with severe or fatal COVID-19 (**Fig.8** – *black lines*). Similarly, we found a positive correlation between epitopes cross-reactivities measured in HD and the corresponding slopes S calculated from **Figs. 1A and 1B** (**Supplemental Fig. S7A** – *upper graph* for CD8^+^ epitopes and *lower graph* for CD4^+^ epitopes) (P<0.0001). Taken together, these results demonstrate that the cross-reactive SARS-CoV-2 epitopes that recalled the strongest CD4^+^ and CD8^+^ T cell responses in unexposed healthy donors (HD) also recalled the strongest responses in asymptomatic COVID-19 patients and were the most highly associated with better disease outcome.

To better understand the possible underlying causes of the observed T-cell cross-reactivity in HD, we determined which of our SL-CoVs-conserved SARS-CoV-2-derived epitopes were also conserved within the four CCCs strains (β-hCCC-HKU1, β-hCCC-OC43 and α-hCCC-NL63, α-hCCC-229E). Using both Multiple Sequences Alignments (MSA) and the Epitope Conservancy Tool (ECT) algorithms and software, we searched for highly similar and identical CD4^+^ and CD8^+^ T cell epitopes potentially cross-reactive between SARS-CoV-2 and the four CCCs strains (**Supplemental Table 3** and **Supplemental Figs. S5** and **S6**). For this, we determined both the percentages of identity (%id) and the similarity scores (S^s^), as described in *Materials* and *Methods*(63). Of the 16 CD4^+^ epitopes, we found ORF1ab_5019-5033_ epitope was highly conserved (%id ≥ 67%) and highly similar (S^S^ ≥ 0.8) between SARS-CoV-2 and the two strains of β-CCCs (β-CCC-HKU1 and β-CCC-OC43), while ORF1ab6088-6102 epitope was highly conserved between SARS-CoV-2 and both β-CCC-HKU1 and α-CCC-NL63 strains (**Supplemental Fig. S5, Supplemental Tables 3** and **4**). Five out of the 27 CD8^+^ epitopes (ORF1ab_3013-3021_, ORF1ab_6749-6757_, S_958-966_, E_20-28_ and M_52-60_) were highly conserved (% id ≥67%) and highly similar (S^S^ ≥ 0.8) between SARS-CoV-2 and the α-CCCs and/or β-CCCs strains. Specifically, the ORF1ab_3013-3021_ CD8^+^ T cell epitope was highly conserved between SARS-CoV-2 and the two strains of β-CCCs (β-CCC-HKU1 and β-CCC-OC43); the ORF1ab_6749-6757_ epitope was highly conserved between SARS-CoV-2 and all the four strains of CCCs; the S_958-966_ epitope was highly conserved between SARS-CoV-2, the two β-CCCs strains and the α-CCC-NL63 strain; the E_20-28_ epitope was highly conserved between SARS-CoV-2 and the β-CCC-HKU1 strain; and the M52-60 epitope was highly conserved between SARS-CoV-2, the two β-CCCs strains and the α-CCC-229E strain (**Supplemental Fig. S6, Supplemental Tables 3** and **4**). While the E_20-28_ epitope was conserved (%id = 67%) between SARS-CoV-2 and α-CCC-NL63 strain, it was not highly similar with the corresponding NL63 peptide (S^S^ = 0.76). Similarly, while the S_976-984_ epitope was conserved between SARS-CoV-2 and three CCCs strains (%id = 67%) it was not highly similar with the corresponding CCC peptides (β-CCC-HKU1 (S^S^=0.78), β-CCC-OC43 (S^S^=0.78) and α-CCC-NL63 (S^S^ = 0.73)). Finally, while the S2-10 epitope was highly similar between SARS-CoV-2 and α-CCC-NL63 (S^S^ = 0.82) it was not highly identical (id% = 56%) (**Supplemental Tables 3** and **4**).

We next determined whether the SARS-CoV-2 epitopes identified above as sharing high identity and similarity with epitopes in various CCCs were targeted preferentially by the CD4^+^ and CD8^+^ T cell responses of either severely ill COVID-19 patients, or of asymptomatic COVID-19 patients (**Fig. 8B**). By comparing the slopes S (**Fig. 1** and **2**) of the SARS-CoV-2-specific CD4^+^ and CD8^+^ T cell responses toward CD4^+^/CD8^+^ epitopes that have neither identical nor similar related peptides in any of the four CCCs (**Fig. 8B** – *first blank column*) with the slopes of T cell responses specific to CD4^+^/CD8^+^ epitopes that are highly similar and/or identical (conserved) with at least one of the four CCCs (**Fig. 8B** – *second blank column*), we could not find any significant differences. By contrast, SARS-CoV-2 CD4^+^ or CD8^+^ T cell responses targeting epitopes conserved (highly identical and similar) *exclusively* in β-CCCs but not in α-CCCs (i.e., epitopes ORF1ab_5019-5033_ and ORF1ab_3013-3021_) have a significantly lower slope S (*P*=0.04 – **Fig. 8B**). In fact, those two epitopes have their slopes S the closest to 0 among all epitopes (**Supplemental Table 2)** and were not significantly correlated with less disease severity (**Figs. 1 and 2)**. In conclusion, epitopes sharing high identity and similarity *exclusively* with beta CCCs were targeted mainly by severely ill symptomatic patients.

In summary, these results indicate that: (*i*) asymptomatic patients and patients with mild COVID-19 preferentially developed strong IFN-*γ*^+^ CD4^+^ and CD8^+^ T cell responses toward the most cross-reactive SL-CoVs-conserved SARS-CoV-2 epitopes, i.e., the epitopes inducing the highest CD4^+^/CD8^+^ T cell responses in unexposed healthy donors; and (*ii*) compared to asymptomatic COVID-19 patients, the severely ill COVID-19 patients and patients with fatal outcomes developed a SARS-CoV-2-specific CD4^+^ and CD8^+^ T cells response preferentially targeting β-CCCs cross-reactive epitopes. Overall, this suggests that strong pre-existing cross-reactive CD4^+^ and CD8^+^ T cells play a role in shaping SARS-CoV-2-specific protective T cell immunity associated with less severe disease in COVID-19 patients.

## DISCUSSION

In the present study, we report that compared with the (non-vaccinated) asymptomatic COVID-19 patients developing little to no disease, the severely ill symptomatic patients that required admission to an ICU and patients with fatal outcomes exhibited high frequencies of exhausted PD-1^+^TIM3^+^TIGIT^+^CTLA4^+^CD4^+^ and PD-1^+^TIM3^+^TIGIT^+^CTLA4^+^CD8^+^ T cells, and low frequencies of functional SARS-CoV-2-specific IFN-*γ*^+^CD4^+^ and IFN-*γ*^+^CD8^+^ T cells. Interestingly, compared to severely ill COVID-19 patients and to patients with fatal outcomes, the asymptomatic COVID-19 patients were more commonly co-infected with the α-CCCs strains, whereas there was no difference in the prevalence of co-infections with β-CCCs strains in all groups of COVID-19 patients. A recent systematic review and meta-analysis of 95 studies that include 29 million individuals undergoing testing, the pooled percentage of asymptomatic COVID-19 infections was 40.5% among individuals with confirmed SARS-CoV-2 infection(68). While about 20% of COVID-19 patients with confirmed SARS-CoV-2 infection develop severe disease(6), the mechanisms leading to this pathogenesis of COVID-19 are still incompletely understood, though it seems to involve significant immune dysregulations. Severe symptoms have been associated with: (*i*) increased levels of pro-inflammatory cytokines (driven by inflammatory monocytes and neutrophils) (8-10); (*ii*) a general lymphopenia (11-16); and (*iii*) a broad (not SARS-CoV-2-specific T cell exhaustion and/or impaired function (15-32). This was reported for immune cells both in the peripheral compartment (PBMCs) and in the lung and brain of symptomatic patients (13, 69). The association of T cell exhaustion with COVID-19 severity is under debate with one study reporting no clear significant correlation with disease severity(70) (using a small number of patients) while two other reports, also using a small cohort of patients, discounted the link between higher expression exhaustion markers and impaired function of SARS-CoV-2-specific CD4^+^ and CD8^+^ T cells in convalescent patients (71, 72). In contrast to these previous reports, the present study uses larger cohorts of COVID-19 patients with detailed clinical differentiation of symptomatic and asymptomatic patients to demonstrate that high frequencies of phenotypically and functionally exhausted CD4^+^ and CD8^+^ T cells specific to conserved epitopes were associated with severe symptoms in critically ill patients and in patients with a fatal outcome.

We also report here that an early and broad lymphopenia positively correlated with COVID-19 disease severity and mortality, consistent with previous reports(66). Moreover, our study also confirmed previous reports of broad leukocytosis combined with T cell lymphopenia in severe COVID-19 patients and extended those findings by demonstrating the observed T cell lymphopenia was particularly apparent for SARS-CoV-2-specific T cells (11-15, 66, 67). Moreover, compared with asymptomatic COVID-19 patients, severely ill symptomatic patients and patients with fatal outcomes exhibited high frequencies of exhausted PD-1^+^TIM3^+^TIGIT^+^CTLA4^+^CD4^+^ and PD-1^+^TIM3^+^TIGIT^+^CTLA4^+^CD8^+^ T cells. In contrast, the less severe disease in asymptomatic and surviving patients inversely correlated with high frequencies of functional (less exhausted) SARS-CoV-2-specific CD134^+^CD137^+^CD4^+^ and CD134^+^CD137^+^CD8^+^ T cells. Our results also agree with a previous finding that showed increased levels of programmed cell death protein 1 (PD-1) in severe cases compared to those in the non-severe cases(3). In addition, we extend those reports by showing that the exhausted SARS-CoV-2-specific CD4^+^ and CD8^+^ T cells co-express TIM3, TIGIT, and CTLA4 markers of exhaustion, besides PD-1. In addition, we detected low frequencies of SARS-CoV-2-specific IFN-*γ*^+^CD4^+^ and IFN-*γ*^+^CD8^+^ T cells in severely ill symptomatic patients with severe disease or fatal outcomes. This finding confirmed previous reports of impaired cellular functionality in CD4^+^ and CD8^+^ T cells in severe COVID-19 cases along with generally lower interferon gamma (IFN-*γ*) and tumor necrosis factor alpha (TNF-α) production (8, 16, 42, 73). Our data indicates that, early after the onset of disease symptoms, exhaustion of peripheral blood-derived SARS-CoV-2 specific CD4^+^ and CD8^+^ T cells might be a suitable predictor of COVID-19 disease severity.

In the present study and as we previously reported(1), we detected pre-existing cross-reactive CD4^+^ and CD8^+^ T cells specific to many of our SARS-CoV-2 epitopes in 15 healthy donors, who has never been exposed to COVID-19 (**Supplemental Fig. S4**). Data from our group and others (1, 51-53, 55, 74) suggest that the presence of cross-reactive T-cells in uninfected healthy individuals who have never been in contact with SARS-CoV-2 may result, at least partially, from T-cells induced following previous exposure to CCCs infections (37, 50-52, 54) (**Supplemental Figs. S4, S5** and **S6**). Interestingly, compared to the patients with severe COVID-19 the asymptomatic patients presented significantly higher frequencies of co-infections with α-CCCs strains (**Fig. 7**). Conversely, severely ill patients comparatively preferentially responded to SARS-CoV-2 epitopes cross-reacting with β-CCCs solely. Our data suggests that mechanisms of T cell exhaustion may involve prior infections with β-strains of CCCs. It is likely that different repertoires of protective and pathogenic SARS-CoV-2 specific T cells targeting cross-reactive epitopes from structural, non-structural, and regulatory protein antigens are associated with different disease outcomes in COVID-19 patients (73, 75). One cannot rule out, however, that a rapid establishment of α-CCCs-cross-reactive SARS-CoV-2-specific CD4^+^ and CD8^+^ T cell responses resulting from previous exposure to α-CCC strain(s) induced protective T cell immunity that led to less-severe COVID-19 disease. In contrast, β-CCCs-cross-reactive SARS-CoV-2-specific CD4^+^ and CD8^+^ T cell responses resulting from previous exposure to β-CCC strain(s) might lead to immunopathology associated with severe COVID-19 disease. Indeed, we found that concomitant CCCs/SARS-CoV-2 co-infections have different effects on disease severity depending on the CCCs strain: SARS-CoV-2/β-CCCs strain (i.e., HKU1 and OC43) co-infections were correlated with a trend (although not significant) toward more severe COVID-19 disease (**Fig. 7B** and **7C**), whereas SARS-CoV-2/α-CCCs strain (i.e., NL63 and mainly 229E) co-infections significantly correlated with less severe COVID-19 disease. Accordingly, two of our SARS-CoV-2 epitopes that are exclusively conserved in both β-CCCs strains HKU1 and OC43 (sharing high identity and similarity with the corresponding CCCs peptides) did not correlate with less disease severity (and have the lowest S values). β-CCCs share more potential cross-reactive epitopes than α-CCCs, with SARS-CoV-2 itself being in the β genera. With that in mind, and because we observed more exhausted SARS-CoV-2 T cells in severely ill patients, it is likely that not all CCCs genera or strains lead to the same phenotypic pre-existing cross-reactive T cell responses (from highly functional to exhausted), thus impacting COVID-19 severity in a variety of ways (toward less or more symptoms, or no impact at all). Our results do not contradict previous reports highlighting that a prior “original antigenic sin” (OAS) potentially linked to previous CCCs might skew the CCCs-specific SARS-CoV-2 cross-reactive T cells toward an exhausted phenotype (76, 77).

However, in line with a previous report(55), not all SARS-CoV-2 cross-reactive T cells observed in healthy donors (HD) were cross-reactive to CCCs epitopes. SARS-CoV-2 epitopes that have the highest number (within the four CCCs strains) of highly probable cross-reactive CCC peptides (with highest %id and highest similarity scores) are not necessarily the same SARS-CoV-2 epitopes that recalled the strongest T cell responses in HD. For example, the SARS-CoV-2 epitopes ORF1a1350-1365, S1-13, M176-190, and ORF612-26 all recalled strong CD4^+^ T cell responses in HD but have no identical nor similar related peptides in any of the four CCCs. The same observation applies for the CD8^+^ epitopes ORF1ab1675-1683, S1000-1008 and S_1220-1228_ (**Supplemental Fig. S4** and **Supplemental Tables 3 and 4**). Interestingly, eight of the 27 CD8^+^ T cell epitopes (ORF1ab_1675-1683_, ORF1ab_5470-5478_, ORF1ab_6749-6757_, S_2-10_, S_958-966_, S_1220-1228_, E_20-28_ and E_26-34_) shared highly identical sequences (%id equal to 67% to 78%) and six of those also sharing high similarity scores (S^S^≥0.8) with predicted epitopes found in common human pathogens (EBV, Streptococcus pneumoniae, Bordetella pertussis and Corynebacterium diphtheriae) and in widely distributed vaccines (BCG and DTa/wP) (**Supplemental Table 7** and **Supplemental Table 4**). The CD8^+^ T cell responses specific to SARS-CoV-2 epitopes sharing high identity and similarity with DTwP vaccines –but not BCG vaccines– epitopes were significantly more associated with less COVID-19 disease severity (**Supplemental Fig. S7B**). These findings suggests that the pre-existing cross-reactive T cell responses may not be the consequence of a single mechanism, but rather could be shaped by antigens present in various pathogens (including CCCs) and widely administrated vaccines (BCG, DTwP). Indeed, the most functional SARS-CoV-2 conserved CD8^+^ T cell epitopes were highly similar and identical with epitopes from the DTwP vaccine (**Supplemental Table 2** and **7**). These findings are consistent with a previous study that described a correlation between DTwP vaccination and fewer COVID-19 deaths(64). The same hypothesis as above with CCCs can be made regarding other antigenic sources of pre-existing cross-reactive SARS-CoV-2 T cell responses in unexposed healthy individuals, such as allergens, DTw/aP and BCG vaccines and other pathogens such as EBV, Streptococcus pneumoniae, Bordetella pertussis among others (**Supplemental Table 4** and (64, 78, 79)). Even human interactions with various animal coronaviruses might trigger SARS-CoV-2 cross-reactive T cell responses (80-88). Finally, confirming previous reports (25, 35, 36, 75, 89), we found a significant age-dependent and comorbidities-associated susceptibility COVID-19 disease with patient over 60, and those with pre-existing diabetic and hypertension comorbidities, being the most susceptible to severe COVID-19 disease.

The development of the next generation of therapeutics and vaccines will benefit from knowledge of mechanisms at play in the immune dysregulations associated with pathogenesis of COVID-19. Most currently available COVID-19 vaccines (mRNA, nanoparticles, adenoviral vectors) are focused on generating a strong immune response against the surface protein of the virus: Spike (33, 34, 90). By exclusively targeting Spike, such vaccines mainly aim to elicit strong humoral immunity in the form of neutralizing antibodies to block or minimize viral infection (34, 91-93). These vaccines have shown great success in preventing severe COVID-19 (94, 95) and in lowering viral load (96, 97). However, they do not entirely block infection, especially with the newly rising SARS-CoV-2 variants, such as the fast-spreading OMICRON variant (98, 99). Therefore, there are limitations with the current vaccines. First, by applying a strong selection pressure on Spike only, this will likely shape virus evolution towards the appearance of variants with mutations in Spike that can escape vaccine-induced antibody protection (100-102). Second, although the Spike protein seems to generate a T-cell response(34), excluding other viral antigens from the vaccine that could contain immunodominant T cell epitopes (35, 36, 44, 103) may lead to (*i*) a limited repertoire of CD8^+^ T cell responses and (*ii*) generate a CD4^+^ T helper / Tfh response that might not sustain the B-cell memory efficiently (multiple studies underscore the correlation between T and B responses: (25, 35, 36, 75, 89), leading to a reduction in antibody production over time (104, 105). These concerns seem especially relevant in the long term(106) and in the elderly and immunocompromised patients, populations known to be already at risk of developing severe COVID-19 (41, 107, 108). The positive correlation between functional SARS-CoV-2 specific CD4^+^ and CD8^+^ T cells and better disease outcome in asymptomatic COVID-19 patients supports the importance of developing CoVs vaccines that target, not only antibody responses, but also early functional SARS-CoV-2 specific CD4^+^ and CD8^+^ T cell responses. Moreover, these vaccines may benefit from a combination with immune checkpoint blockade to reverse the exhaustion of SARS-CoV-2 specific CD4^+^ and CD8^+^ T cells in individuals who are the most susceptible to severe COVID-19. In addition, it will be important to incorporate select T cell antigens and epitopes associated with less-disease severity and that are conserved across animal and human SL-CoVs. Pre-existing T cells targeting conserved SARS-CoV-2 epitopes that cross-react with α-CCCs, but not β-CCCs, may be important in preventing severe COVID-19 symptoms. We are currently assessing in HLA-A2/DR1 hACE2 triple transgenic mice whether candidate multi-epitope-based pan-SL-CoVs vaccines expressing the best “asymptomatic” epitopes that cross-react with α-CCCs (i.e., excluding epitopes cross-reacting solely with β-CCCs) would induce better protection.

This study has certain limitations worth noting. First, the study did not follow up with the COVID-19 patients at later times during convalescence. Second, since the lymphopenia reported in this study was assessed in the peripheral blood, this may not reflect tissue resident CD4^+^ and CD8^+^ T cells. Also, the severity of COVID-19 disease and the higher mortality risks might be attributed to dysregulation of lung-resident SARS-CoV-2 specific CD4^+^ and CD8^+^ T cells, rather than peripheral blood T cells. Thus, further studies will focus on lung tissue-resident SARS-CoV-2 specific CD4^+^ and CD8^+^ T cells to determine whether they correlate positively with the extent of this lymphopenia. Third, the analyses have not been adjusted retrospectively to previous CCCs infections, due to a lack of pre-COVID-19 samples from our cohort of patients. Fourth, although we measured the early stage of the patients’ CD4^+^ and CD8^+^ SARS-CoV-2-specific T cell responses (blood sampled on average 4.8 days after the appearance of the first reported symptoms – **Table 1** and **Supplemental Table 1**), we cannot be precise about the timing of the patients’ first exposure to SARS-CoV-2. Fifth, the cohort of patients enrolled in this study included 50% of Hispanic population. Nevertheless, our results seem to confirm the hypothesis underscored by others(7) that asymptomatic or mild disease best correlates with the presence of early and more functional (less exhausted) SARS-CoV-2 specific T cell responses against various antigens across the viral proteome. Our findings also extend previous reports by showing that, compared to asymptomatic COVID-19 patients, severely ill symptomatic patients, and patients with fatal outcomes, had more exhausted SARS-CoV-2-speccific CD4^+^ and CD8^+^ T cells that preferentially target cross-reactive epitopes that share high identity and similarity solely with the β-CCCs strains.

In conclusion, this study confirms a broad lymphopenia and reports for the first-time high frequencies of functionally exhausted SARS-CoV-2-specific PD-1^+^TIM3^+^TIGIT^+^CTLA4^+^ CD4^+^ and PD-1^+^TIM3^+^TIGIT^+^CTLA4^+^ CD8^+^ T cells were associated with severe disease in critically ill COVID-19 patients (having often more pre-existing diabetes and hypertension co-morbidities). Moreover, compared to severely ill COVID-19 patients and to patients with fatal outcomes, the (non-vaccinated) asymptomatic COVID-19 patients presented more co-infections with the α-CCCs strains and presented more functional SARS-CoV-2-specific CD4^+^ and CD8^+^ T cells that targeted cross-reactive epitopes from structural, non-structural, and regulatory proteins. Our findings support the critical role of cross-reactive SARS-CoV-2-specific CD4^+^ and CD8^+^ T cells in protection against severe COVID-19 disease and provide a roadmap for the development of next-generation T-cell based, multi-antigen, pan-Coronavirus vaccines capable of conferring cross-strain protection.

## ACKNOWLEDGMENTS

The authors would like to thank Dr. Dale Long from the NIH Tetramer Facility (Emory University, Atlanta, GA) for providing the Tetramers used in this study. We thank UC Irvine Center for Clinical Research (CCR) and Institute for Clinical & Translational Science (ICTS) for providing human blood samples used in this study. A special thanks to Dr. Alessandro Ghigi and Dr. Kai Zheng for providing patients’ clinical information. We also thank those who contributed directly or indirectly to this COVID-19 project: Gavin S. Herbert, Dr. Steven A. Goldstein, Dr. Michael J. Stamos, Dr. Suzanne B. Sandmeyer, Jim Mazzo, Dr. Daniela Bota, Dr. Beverly L. Alger, Dr. Dan Forthal, Christine Dwight, Janice Briggs, Marge Brannon, Beverley Alberola, Jessica Sheldon, Rosie Magallon and Andria Pontello.

---

**<u>Supplemental Figure S1</u>**: Genotyping of HLA class-I and class-II in COVID-19 patients with **various degrees of disease severity.** (**A**) Melting curves of the three PCRs performed on COVID-19 blood samples from our N=147 patients to validate either their HLA-DRB1*01:01^+^ genotype (in green – *n* = 76), their HLA-A*02:01^+^ genotype (in blue – *n* = 55) or their HLA-DRB1*01:01^+^ / HLA-A*02:01^+^ genotype (in orange – *n* = 16). One double negative patient is shown as PCR negative control (in red). To determine the HLA-DRB1*01:01 genotype of a patient, two PCRs (“3a” and “3c”) were used as shown in the figure and one PCR was necessary to determine the HLA-A*02:01 genotype, as described in the Material and Methods.

(**B**) Electrophoresis gel migration of the products (amplicons) of the three PCRs for the control double negative patient and for one patient HLA-A*02:01^+^ (S013), one patient HLA-DRB1*01:01^+^ (S036) and one double positive patient (S076).

**<u>Supplemental Figure S2:</u> Experimental plan and gating strategy:** (**A**) shows experimental plan followed for the flow-cytometry experiments and the ELISpot experiments presented in Figs. 1 to 6, starting with the COVID-19 blood samples collection, patient genotyping, PBMCs extraction and peptide stimulation. (**B**) shows the gating strategy applied when analyzing the flow cytometry data presented in Figs. 3 to 6.

**<u>Supplemental Figure S3:</u> Frequencies of EBV (BMLF-1280-288) specific CD8^+^ T cells in COVID-19 patients with various degrees of disease severity:** (**A**) shows the tetramer staining against EBV BMLF-1280-288 specific CD8^+^ T cells after 48 hours stimulation with the corresponding peptide, in three groups of disease severity: severity 0 (ASYMP – 2 patients), severity 1-2 (mild/moderate – 3 patients) and severity 3-4-5 (severe disease – 3 patients). (**B**) Flow cytometry data showing (across the same three groups of disease severity) co-expression of the exhaustion receptors PD1, TIGIT, TIM-3 and CTLA-4 (*two upper panels*) and the expression of the AIMs CD137/CD134 (*lower panel*) in the BMLF-1280-288 tetramers positive cell population (gated in **A**) after peptide stimulation. For both (**A** and **B**), are representative flow-cytometry dot plots (in *right panels*) and in *left panels* are the associated columns graphs with averages/means of the frequencies of the gated cells. Data are expressed as the mean ± SD. Results were considered statistically significant at *P* ≤ 0.05 (one-way ANOVA).

**<u>Supplemental Figure S4:</u> CD4^+^ and CD8^+^ T cell responses specific to SL-CoVs-conserved SARS-CoV-2-derived epitopes, detected in all COVID-19 patients (regardless of disease severity) and in unexposed Healthy individuals:** Both graphs show IFN-*γ* ELISpot data from COVID-19 patients without disease categories breakdown, compared with ELISpot data from unexposed healthy individuals (HD). The *Upper graph* (related to Fig. 1) shows average SFCs after 72 hours CD4-peptide stimulation of COVID-19+ HLA-A*02:01^+^ patients’ PBMCs (*n* = 71; black bars: SARS-CoV-2 specific CD4^+^ T cell response) or of HD’ PBMCs (*n* = 15; white bars: SARS-CoV-2 cross-reactive CD4^+^ T cell response). Likewise, the *lower graph* (related to Fig. 2) shows average SFCs after 72 hours CD8-peptide stimulation of COVID-19+ HLA-DRB1*01:01^+^ patients’ PBMCs (*n* = 92; black bars: SARS-CoV-2 specific CD8^+^ T cell response) or of HD’ PBMCs (*n* = 15; white bars: SARS-CoV-2 cross-reactive CD8^+^ T cell response). A mean SFCs between 25 and 50 correspond to a medium/intermediate response whereas a strong response is defined for a mean SFCs > 50 per 0.5x10^6^ stimulated PBMCs.

**<u>Supplemental Figure S5 and S6</u>: Best matching sequences of CCCs epitopes with 16 CD4^+^ (Fig. S5) and 27 CD8^+^ (Fig. S6) SARS-CoV-2-derived epitopes:** Matching CCCs peptides were chosen after combining both MSA and ECT analysis (see Materials and Methods, **Supplemental Table 5,** and **Supplemental Table 3**). Each panel in both figures represent the alignment of one SARS-CoV-2 epitope and the four corresponding CCCs best matching peptide sequences. SARS-CoV-2 peptide sequence is set as 100% identity. The Amino Acids color-code was generated with Gecos software (https://gecos.biotite-python.org) using the following parameters: gecos --matrix BLOSUM62 --lmin 60 --lmax 75 –f. As a result, the distance between two Amino Acids in the substitution matrix (BLOSUM62) corresponds to the perceptual visual differences in the color scheme. Similarity score (S^S^) based on such matrix are a good predictive measure of potential cross-reactivity (along with % of peptide identity). S^S^ ≥ 0.80 and %id ≥ 67% are in red. Identity percentages, Similarity scores, conservation and consensus sequences are indicated in both figures for each panel.

**<u>Supplemental Figure S7:</u> Relation between the SARS-CoV-2-crossreactive pre-existing T cell responses in unexposed HD and the magnitude of the correlation between the related epitope-specific T cell responses in SARS-CoV-2 infected patients and the protection against severe COVID-19: (A)** Both graphs represent the correlations between the cross-reactivity of each epitope in HD (i.e., the average number of SARS-CoV-2-specific cross-reactive IFN-*γ*-producing T cells measured by ELISpot in unexposed healthy donors – **Supplemental Fig. S4** – after stimulation with each of the 16 CD4 epitopes, *upper graph*; or the 27 CD8 epitopes, *lower graph*), and the Slope S of the Lines of Best Fit from Fig. 1C (for CD4 epitopes) and Fig. 2C (for CD8 epitopes). S is used as a measure of the magnitude of the correlation between the breadth of each epitope-specific T cell response in patients and the corresponding COVID-19 disease severity (for every epitope: the higher S, the more a strong T cell response toward this epitope is associated to better disease outcome). For both graphs, indicated: the coefficient of determination (R^2^), the Pearson correlation coefficients (R), its associated *P*-value and the equation of the best-fitted line calculated (from linear-regression analysis). (**B**) Each of our SL-CoVs-conserved SARS-CoV-2-derived CD8 epitope, its corresponding S value (slope from Fig. 2C) is shown. Here, each CD8 epitope are categorized according to their similarity and % of identity with peptides found in common vaccines and/or common human pathogens (see **Supplemental Table 7**: high identity is defined for %id>67% and high similarity for a similarity score S^S^≥0.8). Epitopes’ cross-reactivity categories are compared using one-way ANOVA and results were considered statistically significant at *P* ≤ 0.05.

**Supplemental Table 1: Detailed demographic features, age, HLA-genotyping, clinical parameters, onset of symptoms and prevalence of comorbidities of each of the 147 patients enrolled in the study:** This table shows all the detailed information (age, sex, race/ethnicity, length of stay, HLA-genotyping, all the experienced symptoms, symptoms onset and the potential comorbidities…) for each individual patient included in the study. Patients medical record numbers were anonymized by assigning each patient a code as follow: AS## for ASYMP patients and S### for SYMP patient (with # being a digit).

**Supplemental Table 2: Detailed information and listing of the 27 class-I-restricted SL-CoVs-conserved SARS-CoV-2-derived CD8 epitopes and the 16 class-II-restricted SL-CoVs-conserved SARS-CoV-2-derived CD4 epitopes:** Regrouped information from our SL-CoVs-conserved SARS-CoV-2-derived CD4 (upper part) and CD8 (lower part) epitopes, such as: epitopes name/position, SARS-CoV-2 corresponding protein, peptides amino-acid sequence, correlation coefficients R and Slopes S (from **Fig 1C** and **2C**). S is used to assess (for each individual SARS-CoV-2 epitope) the magnitude of the correlation between the breadth of this epitope-specific T cell response measured in SARS-CoV-2 infected patients and the protection against severe COVID-19. The blue/red color code allows to visually compare different correlation magnitudes between the SARS-CoV-2 epitopes. For each SARS-CoV-2 epitopes, significance (*P < 0.05*) of each correlation is also indicated, along with the magnitude of the T cell cross-reactive response measured by IFN-γ ELISpots in HD individuals (**Supplemental Fig. 4**).

**Supplemental Table 3: Best matching peptide sequence between SARS-CoV-2 epitopes and CCCs peptides, with identity percentages and similarity scores:** CCCs peptides sequences, names/positions, Identity percentages and Similarity scores (S^s^) with their related SARS-CoV-2 epitopes are detailed in this Table. Details of the CCCs peptide selection method and similarity scores calculations are in *Materials* and *Methods* section, **Supplemental Tables 5 and 6**. The peptide similarity score “S^s^” calculation use here the BLOSUM62 matrix to compare a pair of peptides (peptide “x” from SARS-CoV-2 and “y” from CCC) and is based on the Sune Frankild et al. methodology. 0 ≤ SS ≤ 1: the closest SS is to 1, the highest is the potential for T cell cross-reactivity response toward the related pair of peptides. We used a threshold of S^S^ = 0.8 to discriminate between highly similar and non-similar peptide. Compared to **Supplemental Table 5** (where the peptide selection is solely based on MSA analysis), peptides that were changed based on Epitope Conservancy Tool (ECT) analysis (**Supplemental Table 6**) are highlighted in beige. Highlighted in yellow: following ECT analysis and compared to **Supplemental Table 5** (MSA analysis), those are new hits of highly identical and/or similar CCC peptides for which either the % of identity is ≥ 67%, or with a Similarity score S^S^ ≥ 0.8.

**Supplemental Table 4: T-cell cross-reactivity potential toward the best matching peptide spanned across the four human common cold coronaviruses (CCCs) proteomes and potential cross-reactive epitopes in other common human pathogens and widely administrated vaccines:** For CCCs potential cross-reactive peptides: values (and corresponding color) reflect the potential of cross-reactivity with a CCC peptide (**Supplemental Table 3**). **0**: low to no potential for an CCC peptide to induce a cross-reactive response toward the corresponding SARS-CoV-2 epitope and vice-versa i.e., %id with the corresponding SARS-CoV-2 epitope < 67% AND similarity score S^S^< 0.8; **0.5**: there is a CCC peptide that may induce a cross-reactive response i.e., %id with the corresponding SARS-CoV-2 epitope ≥ 67% OR similarity score S^S^≥ 0.8; **1**: there is a CCC peptide very likely to induce a cross-reactive response i.e., %id ≥ 67% AND S^S^≥ 0.8.

For identification of potential cross-reactive peptides with our SARS-CoV-2 epitopes in widely administrated vaccines and common human pathogen: details are in **Supplemental Table 7**. *In bold and blue*: contain a peptide with very high potential of cross-reactivity with the SARS-CoV-2 epitope (% id ≥ 78% AND Similarity score S^S^≥ 0.8). *In black (non bolded)*: contain a peptide with high potential of cross-reactivity with the SARS-CoV-2 epitope (78% ≥ %id ≥ 67% AND Similarity score S^S^ < 0.8).

**Supplemental Table 5: Aligned SARS-CoV-2 epitopes with CCCs peptides determined using Multiple Sequences Alignment – details and calculations for Supplemental** Table 3: Corresponding CCC peptides were determined here after proteins sequences alignments of all four homologous CCCs proteins plus the SARS-CoV-2 related one using various Multiple Sequences Alignments algorithms ran in JALVIEW, MEGA11 and M-coffee software (i.e. ClustalO, Kalign3 and M-coffee -the latter computing alignments by combining a collection of Multiple Alignments from a Library constituted with the following algorithms: T-Coffee, PCMA, MAFFT, ClustalW, Dialigntx, POA, MUSCLE, and Probcons). Results were also confirmed with global and local Pairwise alignments (Needle and Water algorithms ran in Biopython). In case of different results obtained with the various algorithms, the epitope sequence with the highest BLOSUM62-sum score compared to the SARS-CoV-2 epitope set as reference was chosen. For each pair of SARS-CoV-2-epitope / CCCs corresponding peptide, % of identity and similarity score were calculated.

**Supplemental Table 6: Matching epitopes between SARS-CoV-2 and CCCs determined using Epitope Conservancy Tool (ECT) analysis – details and calculations for Supplemental** Table 3: For each one of our 16 CD4^+^ and 27 CD8^+^ SARS-CoV-2 epitopes, we ran the ECT against the entire proteomes of each CCCs. All the CCCs peptides from the top query – i.e., with the highest % of identity – are reported in this table.

**Supplemental Table 7: Analysis of potential SARS-CoV-2 cross-reactive epitopes in other non-coronavirus common pathogens and widely distributed vaccines – details for Supplemental** Table 4: Query performed on the data gathered from “Potential Cross-Reactive Immunity to SARS-CoV-2 From Common Human Pathogens and Vaccines” by Pedro A. Reche in Frontier Immunol. Only the peptides sharing a % of identity ≥ 67% with the corresponding SARS-CoV-2 epitope were extracted and reported in this table and in **Supplemental Table 4**.

## REFERENCES

1. Prakash S, Srivastava R, Coulon PG, Dhanushkodi NR, Chentoufi AA, Tifrea DF, et al. Genome-Wide B Cell, CD4(+), and CD8(+) T Cell Epitopes That Are Highly Conserved between Human and Animal Coronaviruses, Identified from SARS-CoV-2 as Targets for Preemptive Pan-Coronavirus Vaccines. J Immunol. 2021;206(11):2566-82.

2. Ye ZW, Yuan S, Yuen KS, Fung SY, Chan CP, and Jin DY. Zoonotic origins of human coronaviruses. Int J Biol Sci. 2020;16(10):1686–97.

3. Rahimi G, Rahimi B, Panahi M, Abkhiz S, Saraygord-Afshari N, Milani M, et al. An overview of Betacoronaviruses-associated severe respiratory syndromes, focusing on sex-type-specific immune responses. Int Immunopharmacol. 2021;92:107365.

4. Frutos R, Serra-Cobo J, Pinault L, Lopez Roig M, and Devaux CA. Emergence of Bat-Related Betacoronaviruses: Hazard and Risks. Frontiers in microbiology. 2021;12:591535.

5. Cele S, Jackson L, Khan K, Khoury DS, Moyo-Gwete T, Tegally H, et al. SARS-CoV-2 Omicron has extensive but incomplete escape of Pfizer BNT162b2 elicited neutralization and requires ACE2 for infection. medRxiv. 2021.

6. Wu Z, and McGoogan JM. Characteristics of and important lessons from the coronavirus disease 2019 (COVID-19) outbreak in China: summary of a report of 72 314 cases from the Chinese Center for Disease Control and Prevention. Jama. 2020;323(13):1239–42.

7. Bertoletti A, Le Bert N, Qui M, and Tan AT. SARS-CoV-2-specific T cells in infection and vaccination. Cellular & molecular immunology. 2021;18(10):2307–12.

8. Chen G, Wu D, Guo W, Cao Y, Huang D, Wang H, et al. Clinical and immunological features of severe and moderate coronavirus disease 2019. J Clin Invest. 2020;130(5):2620–9.

9. Zhou Y, Fu B, Zheng X, Wang D, Zhao C, qi Y, et al. Pathogenic T cells and inflammatory monocytes incite inflammatory storm in severe COVID-19 patients. Natl Sci Rev. 2020:nwaa041.

10. Cavalcante-Silva LHA, Carvalho DCM, Lima É A, Galvão J, da Silva JSF, Sales-Neto JM, et al. Neutrophils and COVID-19: The road so far. Int Immunopharmacol. 2021;90:107233.

11. Xu B, Fan CY, Wang AL, Zou YL, Yu YH, He C, et al. Suppressed T cell-mediated immunity in patients with COVID-19: A clinical retrospective study in Wuhan, China. J Infect. 2020;81(1):e51–e60.

12. Qin C, Zhou L, Hu Z, Zhang S, Yang S, Tao Y, et al. Dysregulation of Immune Response in Patients With Coronavirus 2019 (COVID-19) in Wuhan, China. Clin Infect Dis. 2020;71(15):762–8.

13. Zhang B, Yue D, Wang Y, Wang F, Wu S, and Hou H. The dynamics of immune response in COVID-19 patients with different illness severity. J Med Virol. 2021;93(2):1070–7.

14. Adamo S, Chevrier S, Cervia C, Zurbuchen Y, Raeber ME, Yang L, et al. Profound dysregulation of T cell homeostasis and function in patients with severe COVID-19. Allergy. 2021;76(9):2866–81.

15. Mahmoodpoor A, Hosseini M, Soltani-Zangbar S, Sanaie S, Aghebati-Maleki L, Saghaleini SH, et al. Reduction and exhausted features of T lymphocytes under serological changes, and prognostic factors in COVID-19 progression. Mol Immunol. 2021;138:121–7.

16. Zeng Q, Li YZ, Dong SY, Chen ZT, Gao XY, Zhang H, et al. Dynamic SARS-CoV-2-Specific Immunity in Critically Ill Patients With Hypertension. Front Immunol. 2020;11:596684.

17. Zheng H-Y, Zhang M, Yang C-X, Zhang N, Wang X-C, Yang X-P, et al. Elevated exhaustion levels and reduced functional diversity of T cells in peripheral blood may predict severe progression in COVID-19 patients. Cellular & molecular immunology. 2020;17(5):541–3.

18. Li M, Guo W, Dong Y, Wang X, Dai D, Liu X, et al. Elevated Exhaustion Levels of NK and CD8+ T Cells as Indicators for Progression and Prognosis of COVID-19 Disease. Frontiers in immunology. 2020;11(2681).

19. Diao B, Wang C, Tan Y, Chen X, Liu Y, Ning L, et al. Reduction and functional exhaustion of T cells in patients with coronavirus disease 2019 (COVID-19). Frontiers in immunology. 2020;11:827.

20. Mazzoni A, Salvati L, Maggi L, Capone M, Vanni A, Spinicci M, et al. Impaired immune cell cytotoxicity in severe COVID-19 is IL-6 dependent. J Clin Invest. 2020;130(9):4694–703.

21. Moon C. Fighting COVID-19 exhausts T cells. Nature Reviews Immunology. 2020;20(5):277-.

22. Shahbazi M, Moulana Z, Sepidarkish M, Bagherzadeh M, Rezanejad M, Mirzakhani M, et al. Pronounce expression of Tim-3 and CD39 but not PD1 defines CD8 T cells in critical Covid-19 patients. Microb Pathog. 2021;153:104779.

23. Vigón L, Fuertes D, García-Pérez J, Torres M, Rodríguez-Mora S, Mateos E, et al. Impaired Cytotoxic Response in PBMCs From Patients With COVID-19 Admitted to the ICU: Biomarkers to Predict Disease Severity. Front Immunol. 2021;12:665329.

24. Arcanjo A, Pinto KG, Logullo J, Leite PEC, Menezes CCB, Freire-de-Lima L, et al. Critically ill COVID-19 patients exhibit hyperactive cytokine responses associated with effector exhausted senescent T cells in acute infection. The Journal of infectious diseases. 2021.

25. Hou H, Zhang Y, Tang G, Luo Y, Liu W, Cheng C, et al. Immunologic memory to SARS-CoV-2 in convalescent COVID-19 patients at 1 year postinfection. J Allergy Clin Immunol. 2021.

26. Melms JC, Biermann J, Huang H, Wang Y, Nair A, Tagore S, et al. A molecular single-cell lung atlas of lethal COVID-19. Nature. 2021;595(7865):114-9.

27. Awadasseid A, Yin Q, Wu Y, and Zhang W. Potential protective role of the anti-PD-1 blockade against SARS-CoV-2 infection. Biomed Pharmacother. 2021;142:111957.

28. Rendeiro AF, Casano J, Vorkas CK, Singh H, Morales A, DeSimone RA, et al. Profiling of immune dysfunction in COVID-19 patients allows early prediction of disease progression. Life Sci Alliance. 2020;4(2):e202000955.

29. Kreutmair S, Unger S, Núñez NG, Ingelfinger F, Alberti C, De Feo D, et al. Distinct immunological signatures discriminate severe COVID-19 from non-SARS-CoV-2-driven critical pneumonia. Immunity. 2021;54(7):1578–93.e5.

30. Janssen NAF, Grondman I, de Nooijer AH, Boahen CK, Koeken V, Matzaraki V, et al. Dysregulated Innate and Adaptive Immune Responses Discriminate Disease Severity in COVID-19. The Journal of infectious diseases. 2021;223(8):1322–33.

31. Modabber Z, Shahbazi M, Akbari R, Bagherzadeh M, Firouzjahi A, and Mohammadnia-Afrouzi M. TIM-3 as a potential exhaustion marker in CD4(+) T cells of COVID-19 patients. Immun Inflamm Dis. 2021.

32. Heming M, Li X, Räuber S, Mausberg AK, Börsch AL, Hartlehnert M, et al. Neurological Manifestations of COVID-19 Feature T Cell Exhaustion and Dedifferentiated Monocytes in Cerebrospinal Fluid. Immunity. 2021;54(1):164–75.e6.

33. Sadarangani M, Marchant A, and Kollmann TR. Immunological mechanisms of vaccine-induced protection against COVID-19 in humans. Nat Rev Immunol. 2021;21(8):475–84.

34. Sahin U, Muik A, Derhovanessian E, Vogler I, Kranz LM, Vormehr M, et al. COVID-19 vaccine BNT162b1 elicits human antibody and T(H)1 T cell responses. Nature. 2020;586(7830):594-9.

35. Brand I, Gilberg L, Bruger J, Gari M, Wieser A, Eser TM, et al. Broad T Cell Targeting of Structural Proteins After SARS-CoV-2 Infection: High Throughput Assessment of T Cell Reactivity Using an Automated Interferon Gamma Release Assay. Front Immunol. 2021;12:688436.

36. Grifoni A, Weiskopf D, Ramirez SI, Mateus J, Dan JM, Moderbacher CR, et al. Targets of T Cell Responses to SARS-CoV-2 Coronavirus in Humans with COVID-19 Disease and Unexposed Individuals. Cell. 2020;181(7):1489–501.e15.

37. Nelde A, Bilich T, Heitmann JS, Maringer Y, Salih HR, Roerden M, et al. SARS-CoV-2-derived peptides define heterologous and COVID-19-induced T cell recognition. Nat Immunol. 2021;22(1):74–85.

38. McMahan K, Yu J, Mercado NB, Loos C, Tostanoski LH, Chandrashekar A, et al. Correlates of protection against SARS-CoV-2 in rhesus macaques. Nature. 2021;590(7847):630-4.

39. Rydyznski Moderbacher C, Ramirez SI, Dan JM, Grifoni A, Hastie KM, Weiskopf D, et al. Antigen-Specific Adaptive Immunity to SARS-CoV-2 in Acute COVID-19 and Associations with Age and Disease Severity. Cell. 2020;183(4):996–1012.e19.

40. Tan AT, Linster M, Tan CW, Le Bert N, Chia WN, Kunasegaran K, et al. Early induction of functional SARS-CoV-2-specific T cells associates with rapid viral clearance and mild disease in COVID-19 patients. Cell reports. 2021;34(6):108728.

41. Bange EM, Han NA, Wileyto P, Kim JY, Gouma S, Robinson J, et al. CD8(+) T cells contribute to survival in patients with COVID-19 and hematologic cancer. Nat Med. 2021;27(7):1280–9.

42. Le Bert N, Clapham HE, Tan AT, Chia WN, Tham CYL, Lim JM, et al. Highly functional virus-specific cellular immune response in asymptomatic SARS-CoV-2 infection. J Exp Med. 2021;218(5).

43. Anft M, Paniskaki K, Blazquez-Navarro A, Doevelaar A, Seibert FS, Hoelzer B, et al. COVID-19 progression is potentially driven by T cell immunopathogenesis. medRxiv. 2020.

44. Saini SK, Hersby DS, Tamhane T, Povlsen HR, Amaya Hernandez SP, Nielsen M, et al. SARS-CoV-2 genome-wide T cell epitope mapping reveals immunodominance and substantial CD8(+) T cell activation in COVID-19 patients. Sci Immunol. 2021;6(58).

45. Schub D, Klemis V, Schneitler S, Mihm J, Lepper PM, Wilkens H, et al. High levels of SARS-CoV-2-specific T cells with restricted functionality in severe courses of COVID-19. JCI insight. 2020;5(20):e142167.

46. Thieme CJ, Anft M, Paniskaki K, Blazquez-Navarro A, Doevelaar A, Seibert FS, et al. Robust T Cell Response Toward Spike, Membrane, and Nucleocapsid SARS-CoV-2 Proteins Is Not Associated with Recovery in Critical COVID-19 Patients. Cell Rep Med. 2020;1(6):100092.

47. Giménez E, Albert E, Torres I, Remigia MJ, Alcaraz MJ, Galindo MJ, et al. SARS-CoV-2-reactive interferon-γ-producing CD8+ T cells in patients hospitalized with coronavirus disease 2019. J Med Virol. 2021;93(1):375–82.

48. Nielsen SS, Vibholm LK, Monrad I, Olesen R, Frattari GS, Pahus MH, et al. SARS-CoV-2 elicits robust adaptive immune responses regardless of disease severity. EBioMedicine. 2021;68:103410.

49. Echeverría G, Guevara Á, Coloma J, Ruiz AM, Vasquez MM, Tejera E, et al. Pre-existing T-cell immunity to SARS-CoV-2 in unexposed healthy controls in Ecuador, as detected with a COVID-19 Interferon-Gamma Release Assay. International journal of infectious diseases : IJID : official publication of the International Society for Infectious Diseases. 2021;105:21–5.

50. Sette A, and Crotty S. Pre-existing immunity to SARS-CoV-2: the knowns and unknowns. Nat Rev Immunol. 2020;20(8):457–8.

51. Le Bert N, Tan AT, Kunasegaran K, Tham CYL, Hafezi M, Chia A, et al. SARS-CoV-2-specific T cell immunity in cases of COVID-19 and SARS, and uninfected controls. Nature. 2020;584(7821):457-62.

52. Mateus J, Grifoni A, Tarke A, Sidney J, Ramirez SI, Dan JM, et al. Selective and cross-reactive SARS-CoV-2 T cell epitopes in unexposed humans. Science. 2020;370(6512):89-94.

53. Braun J, Loyal L, Frentsch M, Wendisch D, Georg P, Kurth F, et al. SARS-CoV-2-reactive T cells in healthy donors and patients with COVID-19. Nature. 2020;587(7833):270-4.

54. Stefano GB, and Kream RM. Convalescent Memory T Cell Immunity in Individuals with Mild or Asymptomatic SARS-CoV-2 Infection May Result from an Evolutionarily Adapted Immune Response to Coronavirus and the ’Common Cold’. Med Sci Monit. 2020;26:e929789.

55. Tan CCS, Owen CJ, Tham CYL, Bertoletti A, van Dorp L, and Balloux F. Pre-existing T cell-mediated cross-reactivity to SARS-CoV-2 cannot solely be explained by prior exposure to endemic human coronaviruses. Infect Genet Evol. 2021;95:105075.

56. Olerup O, and Zetterquist H. HLA-DRB1*01 subtyping by allele-specific PCR amplification: a sensitive, specific and rapid technique. Tissue antigens. 1991;37(5):197–204.

57. Gatz SA, Pohla H, and Schendel DJ. A PCR-SSP method to specifically select HLA-A*0201 individuals for immunotherapeutic studies. Tissue Antigens. 2000;55(6):532–47.

58. Herr W, Ranieri E, Gambotto A, Kierstead LS, Amoscato AA, Gesualdo L, et al. Identification of naturally processed and HLA-presented Epstein-Barr virus peptides recognized by CD4(+) or CD8(+) T lymphocytes from human blood. Proc Natl Acad Sci U S A. 1999;96(21):12033–8.

59. Dolton G, Tungatt K, Lloyd A, Bianchi V, Theaker SM, Trimby A, et al. More tricks with tetramers: a practical guide to staining T cells with peptide-MHC multimers. Immunology. 2015;146(1):11–22.

60. Kuypers J, Martin ET, Heugel J, Wright N, Morrow R, and Englund JA. Clinical disease in children associated with newly described coronavirus subtypes. Pediatrics. 2007;119(1):e70–6.

61. Gunson RN, Collins TC, and Carman WF. Real-time RT-PCR detection of 12 respiratory viral infections in four triplex reactions. J Clin Virol. 2005;33(4):341–4.

62. Cui L-J, Zhang C, Zhang T, Lu R-J, Xie Z-D, Zhang L-L, et al. Human Coronaviruses HCoV-NL63 and HCoV-HKU1 in Hospitalized Children with Acute Respiratory Infections in Beijing, China. Adv Virol. 2011;2011:129134-.

63. Frankild S, De Boer RJ, Lund O, Nielsen M, and Kesmir C. Amino acid similarity accounts for T cell cross-reactivity and for “holes” in the T cell repertoire. PloS one. 2008;3(3):e1831.

64. Reche PA. Potential Cross-Reactive Immunity to SARS-CoV-2 From Common Human Pathogens and Vaccines. Front Immunol. 2020;11:586984.

65. Mukaka MM. Statistics corner: A guide to appropriate use of correlation coefficient in medical research. Malawi Med J. 2012;24(3):69–71.

66. Huang G, Kovalic AJ, and Graber CJ. Prognostic Value of Leukocytosis and Lymphopenia for Coronavirus Disease Severity. Emerg Infect Dis. 2020;26(8):1839–41.

67. Yang B, Chang X, Huang J, Pan W, Si Z, Zhang C, et al. The role of IL-6/lymphocyte ratio in the peripheral blood of severe patients with COVID-19. International immunopharmacology. 2021;97:107569.

68. Ma Q, Liu J, Liu Q, Kang L, Liu R, Jing W, et al. Global Percentage of Asymptomatic SARS-CoV-2 Infections Among the Tested Population and Individuals With Confirmed COVID-19 Diagnosis: A Systematic Review and Meta-analysis. JAMA Netw Open. 2021;4(12):e2137257.

69. Wang N, Vuerich M, Kalbasi A, Graham JJ, Csizmadia E, Manickas-Hill ZJ, et al. Limited TCR repertoire and ENTPD1 dysregulation mark late-stage COVID-19. iScience. 2021;24(10):103205.

70. Wilk AJ, Rustagi A, Zhao NQ, Roque J, Martínez-Colón GJ, McKechnie JL, et al. A single-cell atlas of the peripheral immune response in patients with severe COVID-19. Nature medicine. 2020;26(7):1070–6.

71. Files JK, Boppana S, Perez MD, Sarkar S, Lowman KE, Qin K, et al. Sustained cellular immune dysregulation in individuals recovering from SARS-CoV-2 infection. J Clin Invest. 2021;131(1).

72. Zhang JY, Wang XM, Xing X, Xu Z, Zhang C, Song JW, et al. Single-cell landscape of immunological responses in patients with COVID-19. Nat Immunol. 2020;21(9):1107–18.

73. Sattler A, Angermair S, Stockmann H, Heim KM, Khadzhynov D, Treskatsch S, et al. SARS-CoV-2-specific T cell responses and correlations with COVID-19 patient predisposition. J Clin Invest. 2020;130(12):6477–89.

74. Sekine T, Perez-Potti A, Rivera-Ballesteros O, Strålin K, Gorin JB, Olsson A, et al. Robust T Cell Immunity in Convalescent Individuals with Asymptomatic or Mild COVID-19. Cell. 2020;183(1):158–68.e14.

75. Peng Y, Mentzer AJ, Liu G, Yao X, Yin Z, Dong D, et al. Broad and strong memory CD4(+) and CD8(+) T cells induced by SARS-CoV-2 in UK convalescent individuals following COVID-19. Nat Immunol. 2020;21(11):1336–45.

76. Aydillo T, Rombauts A, Stadlbauer D, Aslam S, Abelenda-Alonso G, Escalera A, et al. Immunological imprinting of the antibody response in COVID-19 patients. Nat Commun. 2021;12(1):3781.

77. Brazil R. Do childhood colds help the body respond to COVID? Nature. 2021.

78. Balz K, Kaushik A, Chen M, Cemic F, Heger V, Renz H, et al. Homologies between SARS-CoV-2 and allergen proteins may direct T cell-mediated heterologous immune responses. Scientific reports. 2021;11(1):4792.

79. Bagheri N, and Montazeri H. On BCG Vaccine Protection from COVID-19: A Review. SN Compr Clin Med. 2021:1–11.

80. Sánchez CA, Li H, Phelps KL, Zambrana-Torrelio C, Wang LF, Olival KJ, et al. A strategy to assess spillover risk of bat SARS-related coronaviruses in Southeast Asia. medRxiv. 2021.

81. Zhang XM, Herbst W, Kousoulas KG, and Storz J. Biological and genetic characterization of a hemagglutinating coronavirus isolated from a diarrhoeic child. J Med Virol. 1994;44(2):152–61.

82. Wang Z, Yang X, Zhong J, Zhou Y, Tang Z, Zhou H, et al. Exposure to SARS-CoV-2 generates T-cell memory in the absence of a detectable viral infection. Nat Commun. 2021;12(1):1724.

83. Wang N, Li S-Y, Yang X-L, Huang H-M, Zhang Y-J, Guo H, et al. Serological Evidence of Bat SARS-Related Coronavirus Infection in Humans, China. Virologica Sinica. 2018;33(1):104–7.

84. Vlasova AN, Diaz A, Damtie D, Xiu L, Toh T-H, Lee JS-Y, et al. Novel Canine Coronavirus Isolated from a Hospitalized Patient With Pneumonia in East Malaysia. Clinical Infectious Diseases. 2021.

85. Oma VS, Klem T, Tråvén M, Alenius S, Gjerset B, Myrmel M, et al. Temporary carriage of bovine coronavirus and bovine respiratory syncytial virus by fomites and human nasal mucosa after exposure to infected calves. BMC Vet Res. 2018;14(1):22.

86. De Angelis ML, Francescangeli F, Rossi R, Giuliani A, De Maria R, and Zeuner A. Repeated Exposure to Subinfectious Doses of SARS-CoV-2 May Promote T Cell Immunity and Protection against Severe COVID-19. Viruses. 2021;13(6).

87. Zheng M, Zhao X, Zheng S, Chen D, Du P, Li X, et al. Bat SARS-Like WIV1 coronavirus uses the ACE2 of multiple animal species as receptor and evades IFITM3 restriction via TMPRSS2 activation of membrane fusion. Emerging Microbes & Infections. 2020;9(1):1567–79.

88. Lednicky JA, Tagliamonte MS, White SK, Elbadry MA, Alam MM, Stephenson CJ, et al. Emergence of porcine delta-coronavirus pathogenic infections among children in Haiti through independent zoonoses and convergent evolution. medRxiv. 2021:2021.03.19.21253391.

89. Yu KK, Fischinger S, Smith MT, Atyeo C, Cizmeci D, Wolf CR, et al. Comorbid illnesses are associated with altered adaptive immune responses to SARS-CoV-2. JCI Insight. 2021;6(6).

90. Samrat SK, Tharappel AM, Li Z, and Li H. Prospect of SARS-CoV-2 spike protein: Potential role in vaccine and therapeutic development. Virus Res. 2020;288:198141.

91. Vidal SJ, Collier AY, Yu J, McMahan K, Tostanoski LH, Ventura JD, et al. Correlates of Neutralization against SARS-CoV-2 Variants of Concern by Early Pandemic Sera. J Virol. 2021;95(14):e0040421.

92. Papageorgiou AC, and Mohsin I. The SARS-CoV-2 Spike Glycoprotein as a Drug and Vaccine Target: Structural Insights into Its Complexes with ACE2 and Antibodies. Cells. 2020;9(11).

93. Liu L, Wang P, Nair MS, Yu J, Rapp M, Wang Q, et al. Potent neutralizing antibodies against multiple epitopes on SARS-CoV-2 spike. Nature. 2020;584(7821):450-6.

94. Polack FP, Thomas SJ, Kitchin N, Absalon J, Gurtman A, Lockhart S, et al. Safety and Efficacy of the BNT162b2 mRNA Covid-19 Vaccine. The New England journal of medicine. 2020;383(27):2603–15.

95. Baden LR, El Sahly HM, Essink B, Kotloff K, Frey S, Novak R, et al. Efficacy and Safety of the mRNA-1273 SARS-CoV-2 Vaccine. The New England journal of medicine. 2021;384(5):403–16.

96. Levine-Tiefenbrun M, Yelin I, Katz R, Herzel E, Golan Z, Schreiber L, et al. Initial report of decreased SARS-CoV-2 viral load after inoculation with the BNT162b2 vaccine. Nat Med. 2021;27(5):790–2.

97. Ke R, Martinez PP, Smith RL, Gibson LL, Achenbach CJ, McFall S, et al. Longitudinal analysis of SARS-CoV-2 vaccine breakthrough infections reveal limited infectious virus shedding and restricted tissue distribution. medRxiv. 2021:2021.08.30.21262701.

98. Regev-Yochay G, Amit S, Bergwerk M, Lipsitch M, Leshem E, Kahn R, et al. Decreased infectivity following BNT162b2 vaccination: A prospective cohort study in Israel. Lancet Reg Health Eur. 2021;7:100150.

99. Hacisuleyman E, Hale C, Saito Y, Blachere NE, Bergh M, Conlon EG, et al. Vaccine Breakthrough Infections with SARS-CoV-2 Variants. New England Journal of Medicine. 2021;384(23):2212–8.

100. Weisblum Y, Schmidt F, Zhang F, DaSilva J, Poston D, Lorenzi JC, et al. Escape from neutralizing antibodies by SARS-CoV-2 spike protein variants. Elife. 2020;9.

101. Subbarao K. The success of SARS-CoV-2 vaccines and challenges ahead. Cell Host Microbe. 2021;29(7):1111–23.

102. Hoffmann M, Arora P, Groß R, Seidel A, Hörnich BF, Hahn AS, et al. SARS-CoV-2 variants B.1.351 and P.1 escape from neutralizing antibodies. Cell. 2021;184(9):2384-93.e12.

103. Ferretti AP, Kula T, Wang Y, Nguyen DMV, Weinheimer A, Dunlap GS, et al. Unbiased Screens Show CD8(+) T Cells of COVID-19 Patients Recognize Shared Epitopes in SARS-CoV-2 that Largely Reside outside the Spike Protein. Immunity. 2020;53(5):1095–107.e3.

104. Favresse J, Bayart JL, Mullier F, Elsen M, Eucher C, Van Eeckhoudt S, et al. Antibody titres decline 3-month post-vaccination with BNT162b2. Emerg Microbes Infect. 2021;10(1):1495–8.

105. Terpos E, Trougakos IP, Karalis V, Ntanasis-Stathopoulos I, Gumeni S, Apostolakou F, et al. Kinetics of Anti-SARS-CoV-2 Antibody Responses 3 Months Post Complete Vaccination with BNT162b2; A Prospective Study in 283 Health Workers. Cells. 2021;10(8):1942.

106. Tartof SY, Slezak JM, Fischer H, Hong V, Ackerson BK, Ranasinghe ON, et al. Effectiveness of mRNA BNT162b2 COVID-19 vaccine up to 6 months in a large integrated health system in the USA: a retrospective cohort study. The Lancet. 2021.

107. Collier DA, Ferreira IATM, Kotagiri P, Datir RP, Lim EY, Touizer E, et al. Age-related immune response heterogeneity to SARS-CoV-2 vaccine BNT162b2. Nature. 2021;596(7872):417-22.

108. Ameratunga R, Longhurst H, Steele R, Lehnert K, Leung E, Brooks AES, et al. Common Variable Immunodeficiency Disorders, T-Cell Responses to SARS-CoV-2 Vaccines, and the Risk of Chronic COVID-19. The Journal of Allergy and Clinical Immunology: In Practice. 2021.

